# Selection Corrected Statistical Inference for Region Detection with High-throughput Assays

**DOI:** 10.1101/082321

**Authors:** Yuval Benjamini, Jonathan Taylor, Rafael A. Irizarry

**Affiliations:** Department of Statistics, Hebrew University; Department of Statistics, Stanford University; Department of Biostatistics and Computational Biology, Dana Farber Cancer Institute; Department of Biostatistics, Harvard University

## Abstract

Scientists use high-dimensional measurement assays to detect and prioritize regions of strong signal in a spatially organized domain. Examples include finding methylation enriched genomic regions using microarrays and identifying active cortical areas using brain-imaging. The most common procedure for detecting potential regions is to group together neighboring sites where the signal passed a threshold. However, one needs to account for the selection bias induced by this opportunistic procedure to avoid diminishing effects when generalizing to a population. In this paper, we present a model and a method that permit population inference for these detected regions. In particular, we provide non-asymptotic point and confidence interval estimates for mean effect in the region, which account for the local selection bias and the non-stationary covariance that is typical of these data. Such summaries allow researchers to better compare regions of different sizes and different correlation structures. Inference is provided within a conditional one-parameter exponential family for each region, with truncations that match the constraints of selection. A secondary screening-and-adjustment step allows pruning the set of detected regions, while controlling the false-coverage rate for the set of regions that are reported. We illustrate the benefits of the method by applying it to detected genomic regions with differing DNA-methylation rates across tissue types. Our method is shown to provide superior power compared to non-parametric approaches.

## 1 Introduction

Due to the advent of modern measurement technologies, several scientific fields are increasingly relying on data-driven discovery. A prominent example is the success of high-throughput assays such as microarrays and next generation sequencing in biology. While the original application of these technologies depended on predefined biologically relevant measurement units, such as genes or singe nucleotide polymorphisms (SNPs), more recent applications attempt to use data to identify and locate genomic regions of interest. Examples of such application include detection of copy number aberrations (Sebat et al., 2004), transcription binding sites (Zhang et al., 2008), differentially methylated regions (Jaffe et al., 2012c), and active gene regulation elements (Song and Crawford, 2010). A data-driven approach is also common among neuroscientists, who search in imaging and functional imaging data for regions that are affected by variations in cognitive tasks (Friston et al., 1994, Hagler et al., 2006, Woo et al., 2014).

As these technologies mature, the focus of statistical inference shifts from individuals to populations. Instead of searching for regions different from baseline in an individual sample, we instead want to compare two or more populations (cancer versus normal, for example) to locate regions of difference. For population level inference, region detection methodology needs to account for between-individual biological variability, which is often non-stationary, as well as for technical measurement noise. Furthermore, sample size is usually small due to high technology and recruitment costs, so variability in the estimates remains considerable. Searching for small regions of consistent signal within such large noisy assays is therefore prune to false detections, if selection within the noise is not properly accounted for. In contrast, the lost opportunity cost that can result from being overly-conservative is equally concerning. Statistical inference therefore provides a rigorous approach that can aid practitioners make informed decisions regarding resource allocation.

In the context of population inference, the most commonly applied approach is to compute marginal p-values at each site, correct for multiplicity, and then combine contiguous significant sites into regions. For example, publications in the high-profile biology journals have implemented these homegrown ad-hoc analysis pipelines (Kundaje et al., 2015, Becker et al., 2011, Pacis et al., 2015, Lister et al., 2013). However, there is no theoretical justification for extrapolating inferences from the single sites to the region. In particular, using the average of the observed values at the selected sites as an estimate for the region will result in a biased estimate; Kriegeskorte et al. (2009) coined the term *circular inference* for such practices in neuroscience, highlighting the reuse of the same information in the search and in the estimation. Furthermore, the power of such methods to detect regions is limited by the power to detect at the individual site, which are noisier and require a-priori stronger multiplicity corrections compared to regions. For example, a region consisting of several almost-significant sites will be overlooked by such algorithms. Finally, there is no clear way how to prioritize regions of different sizes and different correlation structures without more refined statistical methods. Here we describe a general framework that permits statistical inference for region detection in this context.

Although high-profile genomic publications have mostly ignored it, the statistical literature includes several relevant publications to the challenge of providing valid inferential statements for regions of interest within a large continuous map of statistics. Published methods differ in how they summarize the initial information into a continuous map of statistics: smoothing or convolving the measurements with a pre-specified kernel, forming point-wise Z maps, p-value maps (Pedersen et al., 2012), or likelihood maps (Siegmund et al., 2011, Hansen et al., 2012). Regions of interest can be identified from local maxima, by thresholding(Jaffe et al., 2012c) or by segmentation based on hidden Markov models (Kuan and Chiang, 2012) or likelihood (Zhang and Siegmund, 2012). When the shape of the signal is known, for example in data related to transcription binding sites, a map of scan-statistics can be surveyed for maxima (see Cai and Yuan, 2014, for asymptotic analysis). Due to the large number of measurements this approach assume true signal locations are well separated which permits the employment of multiplicity corrections for individual detected regions (Schwartzman et al., 2011, 2013, Sun et al., 2014). Non-parametric methods can use sample-assignment permutation to simulate a non-parametric null distribution even for non-stationary data from an unknown distribution (Jaffe et al., 2012b, Hayasaka and Nichols, 2003).

However, current methods fail to address two important characteristics of the practical challenge. The first is to form inferential statements regarding the effect size, such as estimates and confidence intervals. In genomics high-throughput data it is well known that unknown confounders can often create many small differences between the groups (Leek et al., 2010). These are not regions of biological interest, yet can reach strong statistical significance in highly powered studies. Estimates of effect size rather than just p-value permit the practitioner to discern biological significance. Technically, the problem of non-null inference requires stronger modeling assumptions and more sophisticated methods for treating nuisance parameters compared to significance testing for a fully specified null. Work on confidence intervals in large processes include Weinstein et al. (2013) for individual points rather than regions, and Sommerfeld et al. (2015) for globally bounding the size of the non-null set. We consider providing estimates of the effect in biologically meaningful units and quantification of the uncertainty in these estimates an important contribution.

The second challenge is non-stationarity: in most genomic signals, both the variance and the auto-correlation change considerably along the genome. This behavior is due to different DNA sequence composition(Bock et al., 2008, Benjamini and Speed, 2012), uneven marker coverage, and other biomedical properties affecting the DNA amplification procedures employed by the relevant experimental protocols (Jaffe et al., 2012a). Non-stationarity makes it harder to compare the observed properties of the signal across regions: a region of *k* adjacent positive sites is more likely to signal a true population difference if probe correlation is small, but more likely to be caused by chance variation if probe correlation is very high.

In this paper, we introduce a comprehensive approach for inference for region detection (IRD) that produces selection-corrected p-values, estimators and confidence intervals for the population effect size. We focus on popular set of selection algorithms that, after preprocessing, apply a threshold-and-merge approach to region detection: the map of statistics is thresholded at a given level, and neighboring sites that pass the threshold are merged together. The key idea is to identify for each potential region its *selection event* – the necessary and sufficient set of conditions on the observed estimate vector that lead to detection of the region. For threshold-and-merge algorithms, the selection event can be described as a set of coordinate-wise truncations. Basing inference on the distribution of a test statistic *conditional on the selection event*, corrects the selection bias (Lee et al., 2013, Fithian et al., 2014). Inference for each region is tailored for the local covariance in the region. When further selection is needed downstream, standard family wise corrections can be applied to the list of detected regions.

The paper is structured as follows. The rest of the introduction describes in more detail the specific genomic signal – DNA-methylation – that will be used to demonstrate our method. In Section 2 we present a model for the data generation and define threshold-and-merge selection. In Section 3 we review the conditional approach to selective-inference, which enables the researcher to analyze each selected region separately by adjusting the tests to the conditional distribution. When the data is approximately multivariate normal, the conditional distribution follows a truncated multivariate normal (TMN, Section 3.3). For an individual region, we describe sampling based tests and interval estimates in Section 4. In Section 5 we return our focus to the set of selected regions in the experiment, discussing a secondary adjustment that is needed if only a subset of the originally detected regions are reported. Sections 6 and 7 evaluate the performance of the method on simulated and measured DNA-methylation data.

### 1.1 Motivating example: differentially methylated regions (DNA)

DNA methylation is a biochemical modification of DNA that does not change the actual sequence and is inherited during mitosis. The process is widely studied because it is thought to play an important role in cell development (Razin and Riggs, 1980) and cancer (Feinberg and Tycko, 2004). The great majority of methylation events occurs when a methyl group attaches to CpG site (a cytosine base followed by a guanine base along the chromosome). These potential methylation sites are depleted in genome (less than 1% compared to GpC’s which are at 4%), and the density in which they appear varies widely (Takai and Jones, 2002). The attachment and detachment processes of the Methyl group are stochastic within each cell and can vary between the cells of the biological sample. Current high-throughput technologies measure the proportion of cells in a biological specimen that are methylated giving a value between 0 and 1 for each measured CpG site. For example, the most widely used product, the Illumina *Infinuim array*, produces these proportion measurement at about 450,000 sites (Bibikova et al., 2011).

Unlike the genomic sequence, methylation differs across different tissues of the same individual, changes with age, environmental impacts, and disease (Robertson, 2005). Of current interest is to associate changes in methylation to biological outcomes such as development and disease. Reported functionally relevant findings have been generally associated with genomic regions rather than single CpG sites (Jaenisch and Bird, 2003, Lister et al., 2009, Aryee et al., 2014) thus our focus on differentially methylated regions (DMRs). Because DNA methylation is also susceptible to several levels of stochastic variability (Hansen et al., 2012) statistical inference is necessary. Figure 1 shows two examples for regions of difference found by comparing samples from human colon and lung. Our method estimates confidence intervals for the difference in each of the regions.

**Figure 1:**
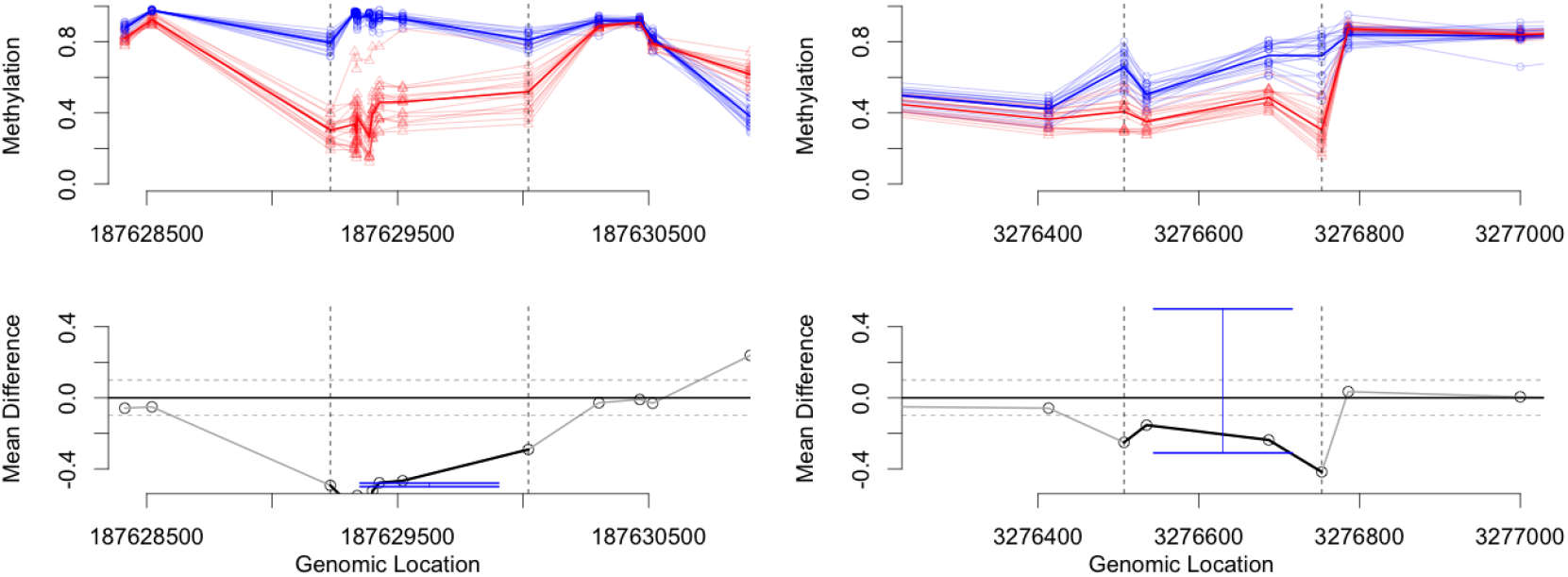
Intervals for differential methylation regions. Confidence intervals for the mean between-tissue different for two candidates regions. Left: A nine-site region (dashed lines) in chromosome 4 and its neighbors plotted by sample (top; tissues are colored and tissue means in bold). The mean-difference (points, bottom-left) considerably exceeds the threshold, and within group variance is small. Hence, the estimated 90% confidence interval (blue) is relatively narrow and the estimate for 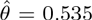, close to the observed mean. Right: A four-site region in chromosome 1 is close to the threshold and the samples do not separate well to groups. Although the estimated mean is 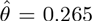, the estimated confidence interval for the mean is not separated from 0. Data was collected by The Cancer Genome Atlas consortium (TCGA).

Inference for DMRs needs to take into account the unknown but variable and often strong local correlation between nearby methylation sites. The propensity for methylation is governed by many local processes, and therefore neighboring methylation sites tend to be have similar methylation proportions. These local influences are thought to be moderated through other local properties such as the DNA attachment to packing proteins, its local chemical properties (Kuan et al., 2010), and the local sequence composition (e.g. the proportion of G and C bases). Therefore, simple correlations models based on the genomic distance between are not sufficient. One of our contributions is an approach that accounts for the non-stationary nature of the data.

## 2 Model for the effects of selection

In the following we introduce a model for the effects of selection on regional descriptors. We begin our description with a population model for individual probes and for the parameter describing the regions, and only then discuss the observed statistic for each regions. We then specify the popular family of threshold-and-merge selection procedures, which we will analyze in this manuscript. Keep in mind that practitioners typically think about these in the opposite direction: observed summaries such as “average observed differences in region” are natural descriptors for the region effect. Selection bias would typically be framed in terms of the expected decay of this observed effect toward zero when the analysis is repeated using the same region but a new set of samples. The population model allows us to estimate and probabilistically bound this decay.

### 2.1 Population model

Suppose the collected data consists of *n* samples of *D* measurements each, *Y*_1_, …,*Y*_*n*_. We model the *i*’th sample as a random process composed of a mean effect that is linear in known covariates and an additive random individual effect. Each sample is annotated by the covariate of interest *X*_*i*_ ∊ *R*, and by a vector of nuisance covariates 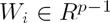. Then the *i*’th observed vector is:

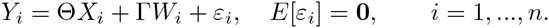

Here 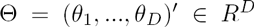 is the fixed process of interest, 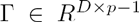 are fixed nuisance processes, and 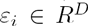 captures both the individual sample effect and any measurement noise. *ε*_*i*_ can further be characterized by a positive-definite covariance matrix 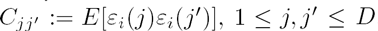. In matrix notation, let 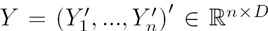 be the matrix of measurements and 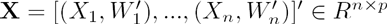 be the design matrix organized so the covariate of interest is in the first column.

For a concrete example of our notation, consider a two-group design comparing samples from two tissue-types. Each sample would be coded in a vector *Y*_*i*_ *∈ R*^*D*^. *X*_*i*_ would code the tissue type, 1 for group A and 0 for group B. *W*_*i*_ can encode such demographic variables as age and gender, as well as a bias term. Then *θ*_*j*_ would code for the mean difference between groups on the *j*’th site. We expect the Θ process 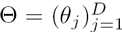 to be almost zero in most sites, and to deviate from zero in short connected regions.

#### Regions of interest

We are interested in detecting and prioritizing regions of the measurement space that are short relative to the size of the genome and for which values in Θ are large in absolute value. A region of interest [ROIs] corresponds to a range *a* : *b* = (*a*, *a* + 1,…, *b*) where 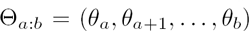 is large (or negatively large) compared to zero. A sufficient representation for an ROI is a triplet *r* = (*a*, *b*, *d*), where *a* ≤ *b ∈* 1,…, *D* are indices and *d ∈* {–, +} represents the direction of deviation from 0. Depending on context, we would usually restrict our analysis to indices *b*, *a* that are *not too far apart*. We code these restrictions into the set 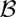 of potential ROIs. Note that 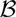 would usually still be very redundant, with many regions that are almost identical; in any run of the algorithm, only a small subset of 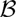 will be selected for estimation.

#### Vector of estimators *Z*

Our inference procedures focus on a vector of point-wise unbiased normal estimators 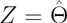. The assumptions for *Z* are:

- Unbiasedness: *E*[*Z*] = Θ.
- Estimable covariance: Σ := Cov(*Z*) is estimable, and the estimate 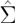 is independent of *Z.*
- Approximate local normality: For indices *a* ≤ *b* that define a potential ROI, meaning (*a*, *b*, +) or (*a*, *b*, −) are in 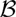, vector *Z*_*a*−1:*b*+1_ is approximately multivariate normal

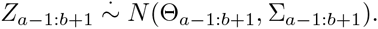

Here, 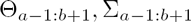 are the 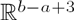 vector and 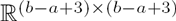 matrix subsets of Θ and Σ respectively.

Specifically, we can take *Z* to be the least-squares estimator for Θ

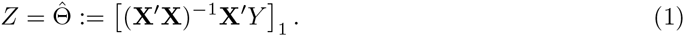

Then, with enough samples compared to covariates (*n* > *p*)^1^, the local covariance *C =* Cov(*ε*_*i*_) is estimable from the linear model residuals, resulting in 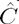. Furthermore, for each 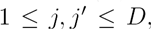, 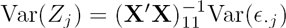 and 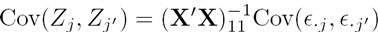. Hence,

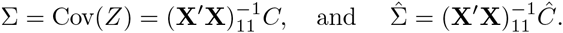

This is extendable to a two-group design where each group is allowed a different covariance.

#### Effect size and population effect size

A commonly used summary for the effect size in a region is the area under the curve (AUC) of the observed process, which is the sum of the estimated effects in the region (Jaffe et al., 2012b). To decouple the region length and the magnitude of the difference, we define the observed effect size as the average rather than the sum of the effect:

##### Definition 1

*The* observed effect size *of region a* : *b is*

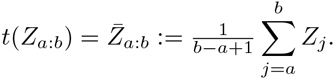

We will associate the observed effect size *t*(*Z*_*a*_:_*b*_) for each potential region *a* : *b* with the population parameter representing its unconditional mean

##### Definition 2

*The* effect size *of region a* : *b is*

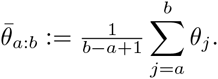

We prefer 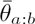 to *AUC* because it decouples two different sources of information: the region length and the effect magnitude. Because correlation varies across different regions, the region length does not correspond monotonically to the amount of independent information. The size of the region, together with the correlation structure, would nevertheless affect the uncertainty of the estimates. Finally note that since the AUC for (*a*, *b*, +) is 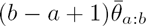, any inference for 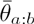 can be easily converted into inference for the AUC. The methods described here are easily extendible for other linear statistics.

### 2.2 Selection

A common way to identify potential ROIs is to screen the map of statistics at some threshold, and then merge sites that passed the screening into regions (Jaffe et al., 2012b, Siegmund et al., 2011, Schwartzman et al., 2013, Woo et al., 2014). This procedure ascertains a minimal biological effect-size in each site, while increasing statistical power to detect regions and controlling the computational effort. After preprocessing and smoothing the responses, these algorithms run a version of the following steps:

1. Produce an unbiased vector of linear estimates 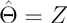.
2. Identify the set of indices exceeding a fixed threshold *c*, {*j* : *Z*_*j*_ > *c*}. This is the *excursion set.*
3. Merge adjacent features that pass the threshold into regions.
4. Filter (or split) regions that are too large.

The procedure is illustrated in Figure 2. The output of such algorithms is a random set of (positive) detected ROIs 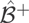 that consists of ROIs (*a*, *b*, +) corresponding to the beginning and end indices of *positive* excursion regions. Step 4 ascertains that 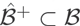.

**Figure 2:**
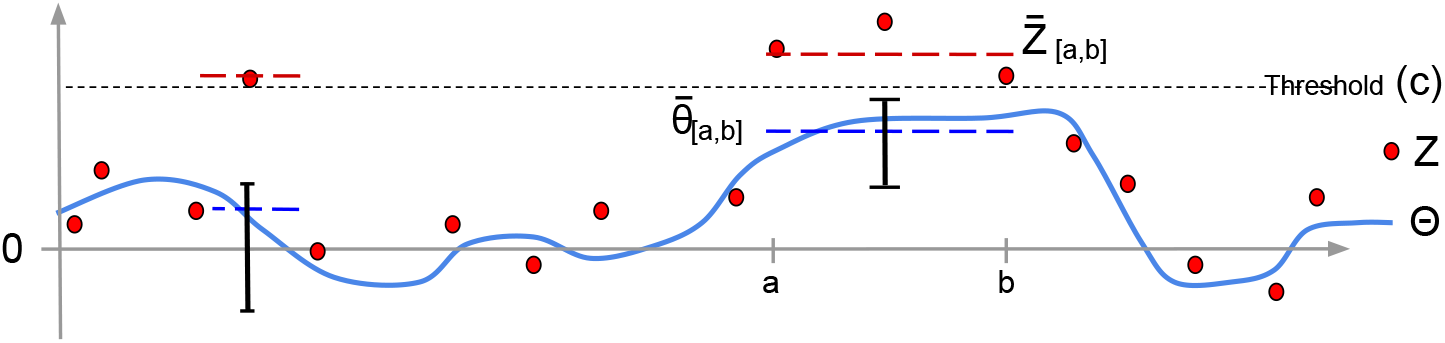
Cartoon of the statistical setup. The parameter vector of interest Θ (solid blue) is unobserved; we observe an unbiased estimate vector Z (full red os). The thresholds (dotted line) are at *c* and −*c*, and the excursion set {*j* : *Z*_*j*_ > *c*} is clustered into two regions. (No regions {*j* : *Z*_*j*_ < −*c*} are shown). Due to this selection, the two parameters to be estimated are 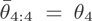 on the left and 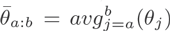, marked with a blue dashed line (here *a* = 8, *b* = 10). The observed effect sizes (red dashed line) are biased because of the selection. Our goal is to form confidence intervals for 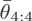 and 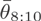.

Comments:

- Typically, we are also interested in the similarly defined set of negative ROIs 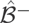. For ease of notation we deal only with 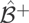, understanding that 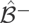 can be analyzed in the same way. It is important to note that each region is selected either to the positive set or to the negative set, and this choice will instruct the hypothesis test.
- Smoothing of the features and local adjustment of the thresholds can both be incorporated within this framework – implicitly changing Θ into a smoothed version.

#### Distribution after selection

It is important to distinguish between inference for predefined regions based on previous biological knowledge, for example exons or transcription start sites, and inference for regions detected with the same data from which we will construct inferential statements. For a *predefined* region, the linear estimator 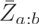 is an unbiased estimator for 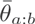, and its uncertainty can be assessed using classical methods. Moreover, 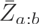 would be approximately normal, so the sampling distribution of 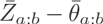 would only depend on a single parameter for the variance. In particular, the correlation between measurements would affect the distribution of 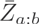 only through this variance parameter 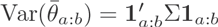, and can be accounted for in studentized intervals.

In contrast, when the ROI (*a*, *b*, +) is detected from the data, 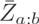 becomes biased (Berk et al., 2013, Fithian et al., 2014, Kriegeskorte et al., 2009). We will tend to observe extreme effect sizes compared to the true population mean. If 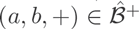, the observed effect size 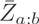 would always be greater than the threshold *c*. Furthermore, the distribution of 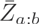 would be right-skewed, so normal-based inference methods are no longer valid.

For non-stationary processes, evaluating the observed regions poses an even greater challenge. Due to variation in the dependence structure, it is no longer straightforward to compare different regions found on the same map. The bias and skewness depend not only on a single index of variance, but rather on the local inter-dependence of *Z* in a neighborhood of *a* : *b*. In Figure 3, we study the effect of correlation on the bias and skewness in a simulated example; changes in correlation affect the bias, the spread and the skewness of the conditional distribution.

**Figure 3:**
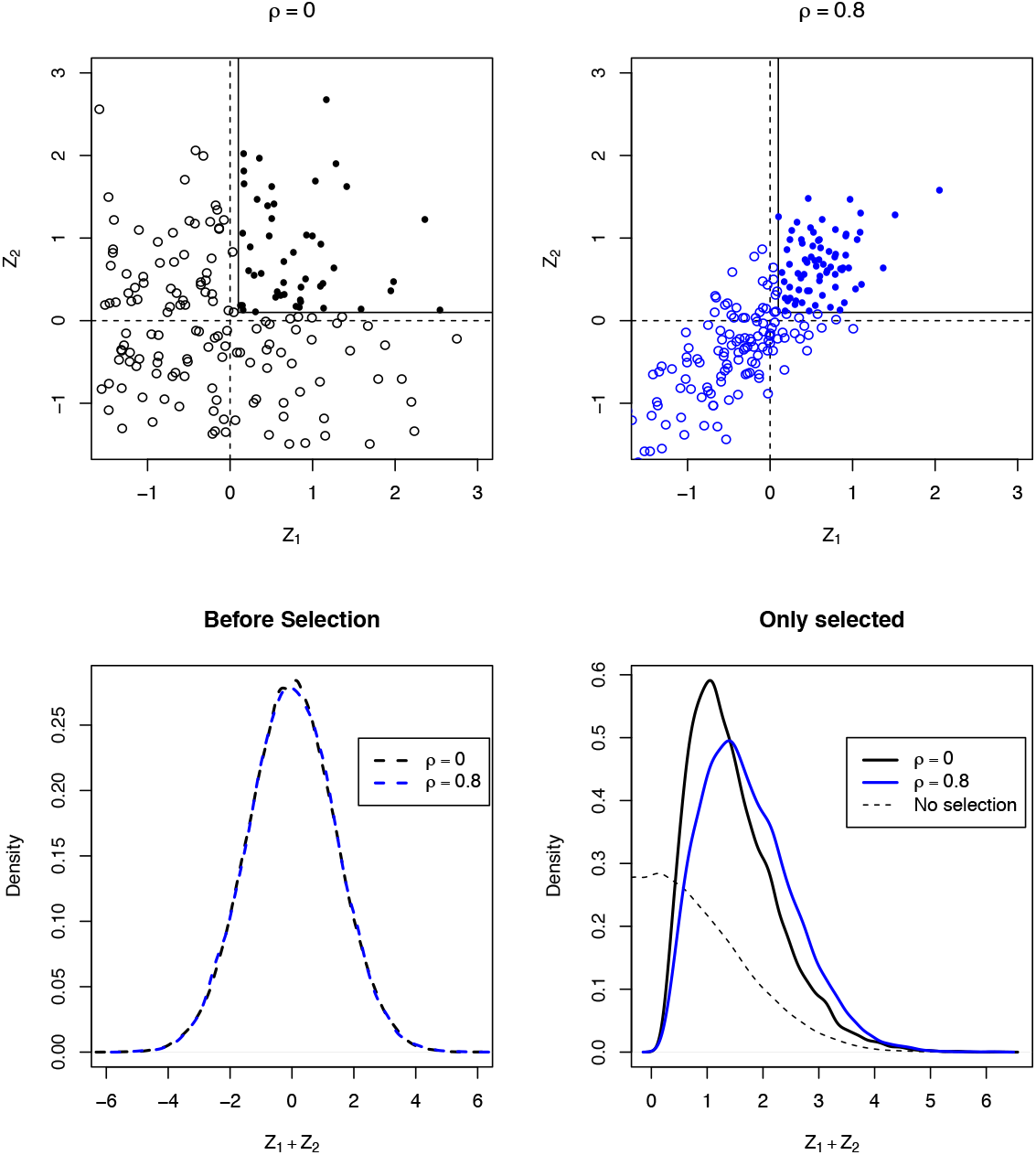
Effects of selection. A simulated example to illustrate the bias and skewness of selected distributions with different correlation parameters. For *D* = 2 and *c* = 0.5, we simulate data from *Z =* (*Z*_1_, *Z*_2_) ~ *N* ((0, 0), (1, *ρ; ρ,* 1)) and consider only cases where the region 1 : 2 was selected. The top plots show the distributions for *ρ* = 0 and *ρ* = 0.8, and the bottom plots show the (rescaled) distribution for the observed effect-size 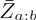 for all data (left) versus the selected data (right). Although the 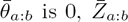 is biased away from 0 (lower right panel); furthermore, each correlation regime shows a different bias and a different distribution. Contrast this with the case of no selection (lower left panel), where there is no bias and no effect of correlation after rescaling.

#### 2.3 Inference goals

We can now restate our goal using the notation. For the set of *K* detected regions 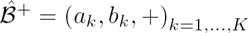, we would like to make valid inferential statements about each 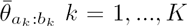 that account for the selection. These include:

1. a hypothesis test for 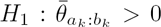 against the null 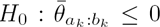, and a high-precision p-value for downstream multiplicity corrections;
2. an estimate for 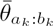;
3. a confidence interval for 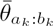.

We would also like to be able to prune the detected set 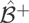 based on the hypotheses tests, while continuing to control for the false coverage statements in the set.

## 3 Conditional approach to selective inference

Because we are only interested in inference for selected ROIs in 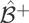, a correction for the selection procedure is needed. Most such corrections are based on evaluating, ahead of selection, the potential family of inferences. In hypothesis testing, this requires, essentially, to evaluate all potential ROIs and transform them into a common distribution (usually by calculating the p-value). Instead, we adapt here a solution proposed for meta-analysis and more recently in the context of regression model selection problems (Lee et al., 2013, Fithian et al., 2014): adjust inference to hold for the conditional distribution of the data given the selection event. First we review the premise of selective inference, and then specify the selection event and selective distribution for region detection.

### 3.1 Selective inference framework

Recall that 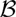 denotes the set of potential ROIs, and the random set 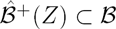 denotes the random set of selected ROIs. Assume that for ROI 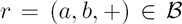 we associate a null hypothesis 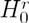 that will be evaluated if and only if 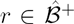. A hypothesis test controls for the *selective error* if it controls for the probability of error given that the test was conducted. Formally, denote by *A*_*r*_ *= A*_(*a,b*,+)_ the event that *r* was selected for 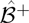. Then:

#### Definition 3 (Control of selective type 1 error, Fithian et al. 2014)

*The hypothesis test φ*_*r*_(*Z*) *∈* {0,1}, *which returns 1 if the null is rejected and 0 otherwise, is said to control the* selective type 1 error *at level α if*

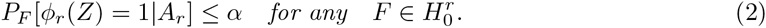

In a frequentist interpretation, the relative long-term frequency of errors in the tests of *r* that are carried out should be controlled at level *α.*

Selective confidence intervals can be defined in a similar manner. Denote by *F* the true distribution of *Z,* so that *F* belongs to a model 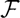. Associate with each *r* a functional *η*_*r*_(*F*) of the true distribution of *Z,* and again let *A*_*r*_ be the event that the confidence interval *I*_*r*_(*Z*) for *η*_*r*_(*F*) is formed. Then *I*_*r*_(*Z*) is a *selective* 1 — *α confidence interval* if:

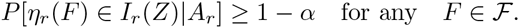

Correcting the tests and intervals to hold over the selection criteria removes most biases that are associated with hypothesis selection. If no additional selection is performed, the selective tests are not susceptible to the “winner’s curse”, whereby the estimates of the selected parameters tend to display over-optimistic results. If each individual test *ϕ*_*r*_ controls selective error at level *α*, then the ratio of mean errors to mean selected is also less than *α*. In a similar manner, the proportion of intervals not covering their parameter (False Coverage Rate, FCR) is controlled at *α* (Fithian et al., 2014, Weinstein et al., 2013). This strong individual criterion allows us to ignore the complicated dependencies between the selection events; as long as the individual inference for the selected ROIs is selection controlled, error is also controlled over the family.

Note however that when the researcher decides to report only a subset of the selected intervals or hypotheses, a secondary multiplicity correction would be required. A likely scenario is reporting only selective intervals that do not cover 0 (Benjamini and Yekutieli, 2005). The set of selective p-values (or selective intervals) behaves like a standard hypothesis family, so usual family-wise or false discovery corrections can be used. We discuss this further in Section 5.

### 3.2 Selection event in region detection

For region detection, we can identify the selection event *A*_*r*_ = *A*_(*a,b*,+)_(*Z*) as a coordinate-wise truncation on coordinates of Z. Selection of (*a*, *b*, +) occurs only if all estimates within the region to exceed the threshold. These are the *internal* conditions:

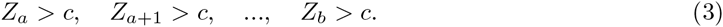

Furthermore, unless *Z*_*a*_ or *Z*_*b*_ are on a boundary, the selection of (*a*, *b*, +) further requires *external* conditions that do not allow the selection of a larger region, meaning:

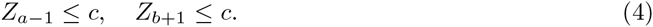

### 3.3 Truncated multivariate normal distribution

Assume the observations are distributed approximately as a multivariate normal distribution *Z* ~ *N*(*Θ,*Σ) with Σ known. According to (2), to form selective tests or intervals based on a statistic *t*_(*a,b*,+)_(*Z*) we need to characterize the conditional distribution of *t*_(*a,b*,+)_(*Z*) given the selection event *A*_(*a,b*,+)_. We will sample the conditional vector to approximate the distribution of the functional.

Conditioning the multivariate normal vector *Z* on the selection event results in coordinate-wise truncated multivariate normal (TMN) vector with density

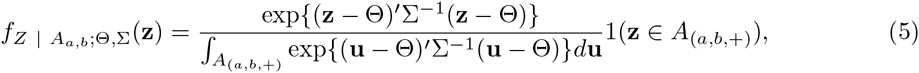

where *A*_(*a,b,*+)_ is seen as a subset of 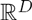. The TMN distribution has been studied in many contexts, including constructing instrumental variables(Lee, 1981), Bayesian inference (Pakman and Paninski, 2014), and lately post selection inference in regressions (Lee et al., 2013). In contrast to the usual multivariate normal, linear functionals of the truncated normal cannot be described analytically (Horrace, 2005). We will therefore resort to Monte Carlo methods for sampling TMN vectors, and empirically estimate the functional distribution from this sample. Naively, one can sample from this distribution using a rejection sampling algorithm: produce samples from the unconditional multivariate normal density of *Z*, and reject samples that do not meet the criteria *A*_(*a,b,*+)_.

In the rest of this paper we assume we have efficient samplers for the TMN distribution. There are many publicly available samplers for this distribution, for example, in the R statistical package, including rejection samplers, Gibbs samplers (Geweke, 1991) and Hamiltonian (Pakman and Paninski, 2014) samplers. We use a Gibbs sampler included in the *selective-inference* package, because it handles well extreme values. Sampling strategies are discussed in Appendix A.

For inferences of a specific index range, it is sufficient to focus on the coordinates of *Z* in the vicinity of *a* : *b*. Coordinates outside the selection range *a −* 1 : *b* + 1 are neither restricted nor influence *t*(*Z*) and can be ignored in the modeling process assuming the detected bumps are sufficiently separated. Instead of modeling and sampling a *D* dimensional TMN, for each selected ROI of length *l* we sample an *l* + 2 vector.

## 4 Inference for the effect size

Here we describe in sampling based inference prodedures. First, we review the case where the null hypothesis is specified by the full mean vector. Next, we propose a mapping from our parameter of interest 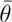 into the mean vector Θ_*a*−1:*b*+1_. For the internal mean, we use a linear mapping 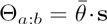 with the *profile* vector s. (In simulations, we find that the conditional distribution of the statistic is not sensitive to small changes in s.) For the external mean *θ*_*a*−1_, *θ*_*b*+1_, we propose a conservative choice of plugging in the observed values 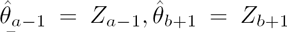. With these choices, the conditional distribution for each value of *theta* is fully specified and sampling based tests can be run. We differ the discussion of efficient sampling strategies to Appendix A.

Notational remarks:

1. Throughout the section we focus on a single selected region *r* = (*a, b*,+). In 4.2 we model only the internal region, so 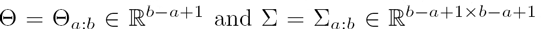. In 4.1, 4.3, we model both external and internal regions, so Θ = Θ_*a*−1:*b*+1_ and Σ = Σ_*a*−1:*b*+1_. We assume here Σ is known, though in practice we will plug in estimates of Σ.
2. When analyzing the positively detected regions 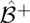, a region *a* : *b* is associated with a single selection event (*a,b*,+). We therefore adopt the shorthand *A*_*a:b*_ = *A*_(*a,b,+*)_. Negatively detected regions would be analyzed separately in a similar manner.
3. Because we are interested primarily in the effects of different mean vectors Θ on the distribution of *Z* and *t*(*Z*) = *Z*_*a:b*_, we use the shorthands

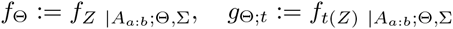

for the TMN density with mean vector Θ and the univariate density of *t*(*Z*) : *Z* ~ *f*_Θ_.

### 4.1 Test for a full mean vector

Consider a vector *Z* that follows a TMN density *f*_Θ_, and suppose we want to test a strong (fully specified) hypothesis 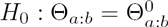 for some 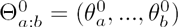 against 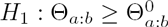. For the test to be powerful against a shift in multiple coordinates, we consider the test

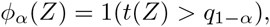

where *t* is a non-negative linear combination 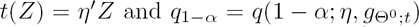 is the 1 – *α* quantile of the distribution of 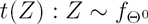. A case in point is the strong null 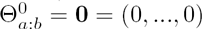. When Σ is known, the null distribution of *t*(*Z*) is fully specified even though its analytic form is unknown. We can therefore sample from 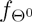, and use this sample to empirically estimate 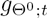 and specifically its 1 − *α* quantile as accurately as we need.

Explicitly, the algorithm would

1. Use a TMN sampler to generate a Monte Carlo sample from 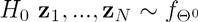.
2. Compute the statistic for each example 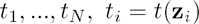.
3. Estimate the 1 – *α* quantile of *t*(*Z*) under *H*_0_ from the 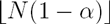 order statistic of *t*_*i*_.

Furthermore, the p-value of the observed vector *z*_*obs*_ can be estimated from the sample as

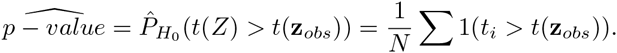

In order to form intervals, we will also need two sided tests. A two sided test can be implemented by setting 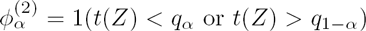 and estimating 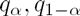 from the Monte Carlo sample. Note that we match *α* level one-sided tests with 1 — 2*α* level intervals so that coverage would agree for the null.

### 4.2 A single parameter family for the mean

Confidence intervals for 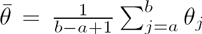 require more care than the null tests, because even after specifying 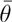 the model has *b* – *a* ancillary degrees of freedom. Approaches to non-parametric inference include plugging-in the maximal-likelihood values of the ancillary parameters for every value of 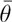 (profile-likelihood, see review in DiCiccio and Romano, 1988), or conditioning on the ancillary directions in the data as in Lockhart et al. (2014), Lee et al. (2013). We take an approach similar to the least favorable one-dimensional exponential family (Efron, 1985), in proposing a linear trajectory from 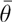 to the mean vector Θ. Figure 4 shows the main steps of our approach.

**Figure 4:**
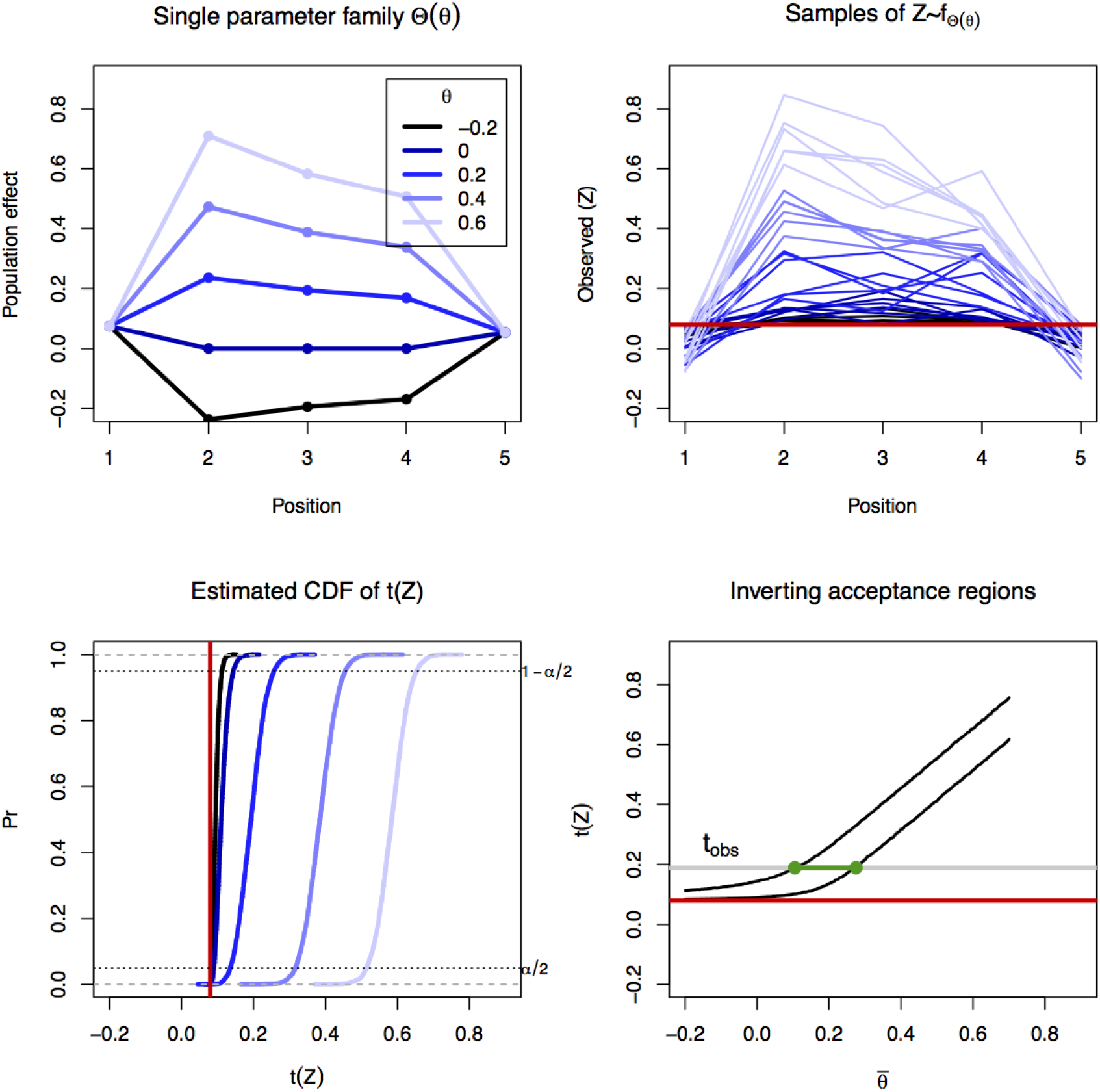
The inference algorithm. For the region 2 : 4 the plots show steps in the inference. The top left displays the unconditional mean vectors Θ_s_(*θ*) for 5 values of *θ*. The profile used is 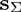. The top right panel displays 6 examples from the Monte Carlo sample of the conditional density 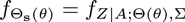, color coded by the value of *θ*. Empirical CDFs are estimated for each value of *θ* (bottom left), and *α*/2,1 – *α*/2 quantiles extracted. The acceptance regions are inverted (bottom right) based on the observed statistic (*t*_*obs*_) to generate a two-sided 1 – *α* interval. Plots based on a simulated region with a true mean effect of 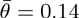 and an observed effect of 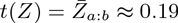. The true (known) covariance of *Z* is used.

We form the confidence interval by inverting a set of tests for the average parameter 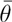. Recall that a random interval *I*(*Z*) is a 1 − 2*α* level confidence interval for 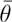 if 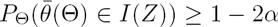 for any Θ. Given a family of 2*α* level two-sided tests for 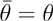, the set of non-rejected parameter-values forms a 1 − 2*α* confidence set (Inversion lemma, Lehmann and Romano, 2005).

The family of tests we use are based on a one-dimensional sub-family of the TMN, produced by the linear (or affine) mapping 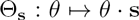, where vector 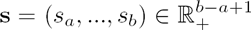 represents the profile of the mean 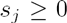 that is scaled linearly by *θ*. For identifiability, we set 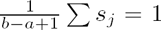. For any value of *θ,* we can define the distribution 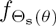 as follows:

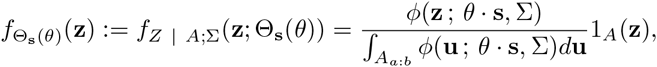

At *θ =* 0, we get the strong null hypothesis Θ = 0. For other values of *θ* we get different TMN distributions. Note that the condition 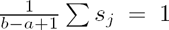 ascertains that 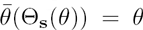, because 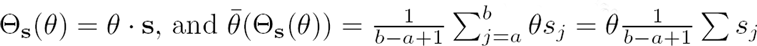.

We derive confidence intervals and point estimators for 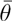 based on this one-parameter family 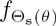. Each value of 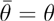 identifies a specific mean vector (panel A), and a conditional distribution for the vector *Z* and the statistic *t*(*Z*) (panels B, C). Therefore, for each value of *θ,* we can construct the two-sided test 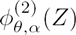 by following the recipe in 4.1. That is, we can sample from 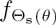, estimate empirically the distribution and quantiles of *t*(*Z*) under this null, and reject if not 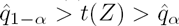. By repeating this process for a fine grid of *θ* values, we can invert the sequence of tests (panel D) and get a high-resolution confidence set or interval *I*.

As a point estimator, we propose using

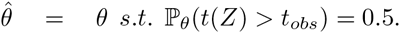

This corresponds to the intersection of the median function 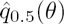 with *t*_*obs*_, and does not require extra calculations when computing the confidence intervals. In Appendix A we show that tests for any value of *θ* can be computed using a single Monte Carlo sample.

For the statistic 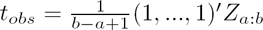, we propose to use the profile:

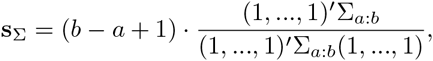

which would vary the mean in each coordinate proportionally to the sum of the columns of the covariance matrix. For the unconditional family parametrized by 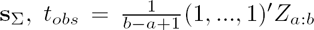 is the natural statistic for the exponential family, as well as the maximum likelihood estimator. After truncation, this is still an exponential family with the same natural statistic, though it is no longer the MLE^2^. An alternative profile is s_*u*_ = (1,…, 1), which induces a uniform increase in the coordinates of the region. In practice, we find that our methods are not sensitive to the particular choice between these two profiles, as seen in Figure 5. Further discussion of the choice of the profile in 4.2.1. Some readers may prefer to skip directly to 4.3.

**Figure 5:**
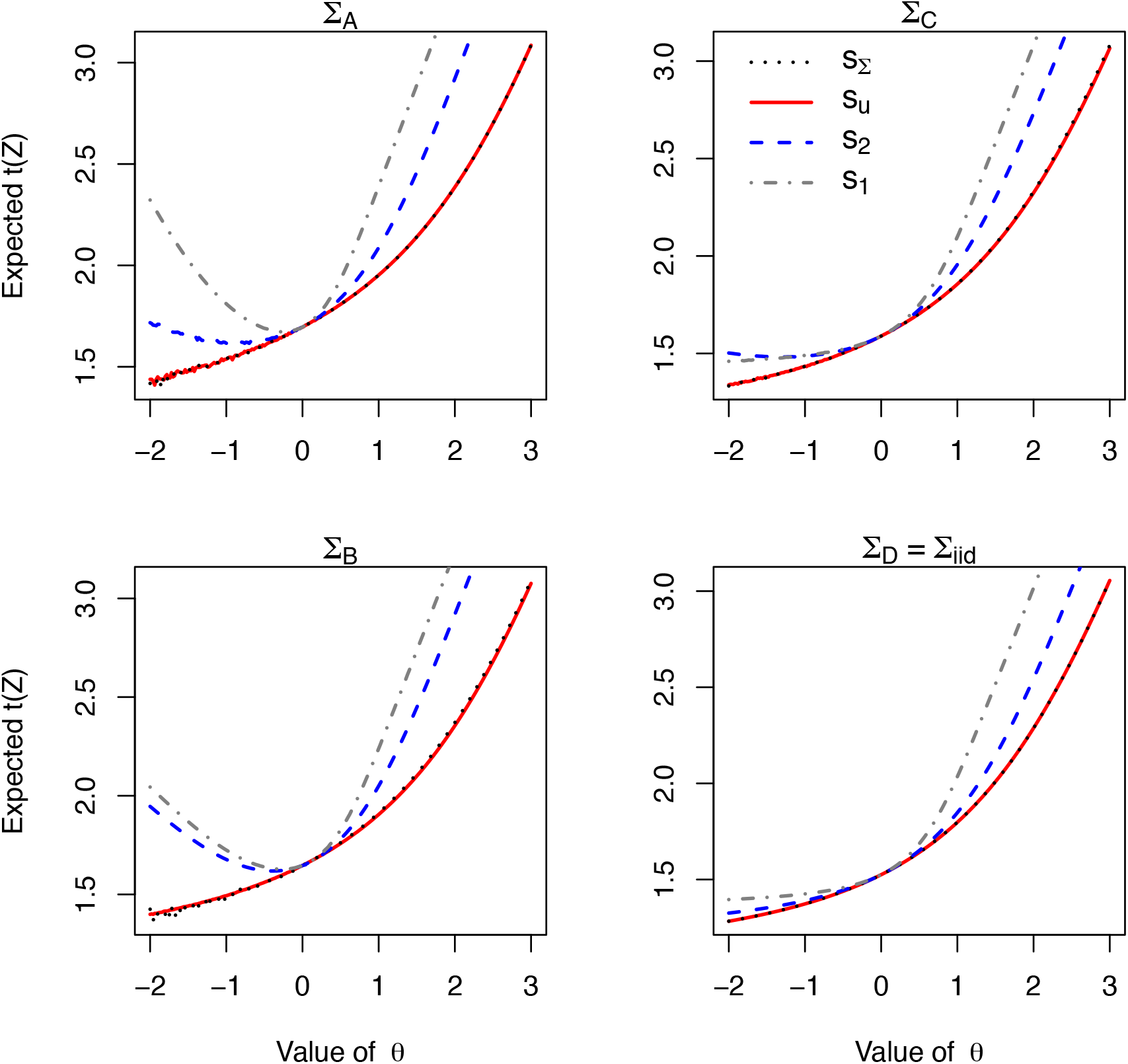
Comparison of profiles vectors and covariances. conditional mean of 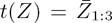 as a function of *θ* for different covariances (panels) and profiles (colors), with threshold *c* = 1. In all panels, we use the following profiles: 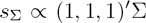 in black, *s*_*u*_ *=* (1, 1, 1) in red, s_2_ = (1.5, 0,1.5) in blue, and s_1_ = (3, 0, 0) in grey. In the top-left and bottom-right panels, *s*_*u*_ ≡ *s*_Σ_). All covariances have unit variance, and the number of correlated variables decrease from Σ_*A*_ (*ρ* = 0.4 between every pair), through Σ_*B*_ (as before but *ρ*_13_ = 0), Σ_*C*_ (*ρ*_12_ = 0) and uncorrelated Σ_*D*_. We observe method is not very sensitive to small differences in the profile, as s_Σ_,s_*u*_ give almost identical curves for Σ_*B*_ and Σ_*C*_. The figure shows that although s_Σ_ ensures 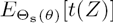 strictly increases with *θ* (monotonicity, Lemma 1), monotonicity is not guaranteed for s_2_ or s_1_. For Σ_*C*_, s_1_ satisfies the conditions of Lemma 2 and displays monotonicity, whereas s_2_ does not. When the covariance is iid, any non-negative profile satisfies Lemma 2. Under all covariances, the curves for s_1_,s_2_ are greater than s_Σ_; this is a potential source for coverage error if s_Σ_ is used.

### 4.2.1 Monotonicity and choice of profile

Although naively we would expect functionals of the distribution of *t*(*Z*) to increase as Θ_s_(*θ*) increases, this monotonicity property does not always hold. Monotonicity is an important prerequisite for the method: measuring *θ* using *t*(*Z*) makes sense only if *t*(*Z*) indeed increases with *θ*. Furthermore, monotonicity guarantees that the acceptance region of the family of tests would be an interval, simplifying the parameter searches. Indeed, for a non-truncated multivariate normal with mean *θ ·* s and s positive, the distribution of *t*(*Z*) is monotone increasing in every coordinate of Θ (and does not depend s). Unfortunately, after conditioning, monotonicity is no longer guaranteed for a general profile vector s; for example, it is not always true that the mean 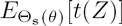 or the quantiles functions will be increasing functions of *θ*. See Figure 5. In the rest of this section we identify conditions for monotonicity: in Lemma 1 we show that for every non-negative covariance, using the profile 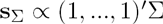 ensures that 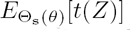 increases in *θ*. This result is tailored for the statistic 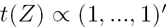. In Lemma 2, we identify a subset of the non-negative covariances and, for each covariance, a set of profiles that ensure monotonicity for any non-negative statistic.

We denote by 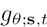 the family of densities parametrized by *θ* of *t*(*Z*)|*A,* where 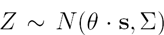. The following lemma proves that for 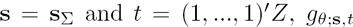 is a monotone likelihood ratio family in *θ*. This is a consequence of identifying *t*(*Z*) as the natural statistic of the family in Fithian et al. (2014).

**Lemma 1** *If g*_*θ*_;_*s,t*_ *denotes the family densities for t*(*Z*) *with the scalar parameter θ as before, then*

1. 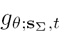 *is a monotone likelihood ratio family.*
2. 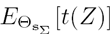 *is an increasing function of θ.*
3. *The confidence set for θ obtained by inverting two sided tests is an interval.*

Properties 2,3 are direct consequences of the monotone likelihood ratio family property. We leave the proof of 1 to the appendix.

The lemma further allows a classical approach to the problem of choosing the statistic after the model. If we choose a specific deviation from 0, e.g. setting the profile to be uniform s_*u*_ = (1,…, 1)ʹ, then the most efficient statistic would be the exponential-family natural statistic 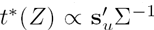. Lemma 1 implies that *t\**(*Z*) would be monotone in *θ.*

A stronger result of monotonicity of 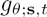 can be derived from properties of the multivariate distribution *f*_Θ_ if the covariance of *Z*_*a,b*_ belongs to a restrictive class of positive-covariance matrices.

#### Definition 4

*An M-matrix is a positive-definite matrix in which all off-diagonal elements are non-positive.*

**Lemma 2** *Suppose* Σ^−1^ *is an M-matrix, and the profile s can be written as a non-negative sum of columns of* Σ, *then g*_*θ*;*s,t*_ *is a monotone likelihood ratio family.*

The condition on *Z* is sufficient to show a strong enough association condition on *Z* (second-order multivariate total positivity property, MTP-2) that continues to hold after conditioning. The lemma is based on theory developed by Rinott and Scarsini (2006). The details of the proof are in the appendix. In Figure 5

## 4.3 Choice of mean for the external constraints

For the distribution *f*_Θ_ to be fully specified, we need to set values for the external mean parameters *θ*_*a*−1_,*θ*_*b*+1_. Although it is possible to arbitrarily assume *θ*_a−1_ = *θ*_*b*+1_ = 0, when the variance of *Z*_*a*−1_ and Z_*b*+1_ (Σ_*a*−1 *a*−1,_ Σ_*b*+1,*b*+1_) are small, this could lead to a bad fit for the data and produce ill-behaved intervals. Scaling the external mean with *θ* also produces artifacts when the *Z*_*a*−1_ or *Z*_*b*+1_ are strongly correlated with (*Z*_*a*_,…, *Z*_*b*_).

We recommend using the plug-estimators for *θ*_*a*−1_, *θ*_*b*+1_ based on the observed values *Z*_*a*−1_, *Z*_*b*+1_:

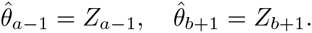

Because selection is often weak for the external parameters, the plug-in estimators seem to do a good job even as Σ decreases.

To summarize, an affine model for the mean is:

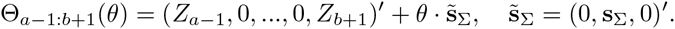

## 4.4 Estimated covariance

When the number of samples is small, the estimation may be sensitive to the covariance that is used. We therefore propose using an inflated estimate of the sample covariance in order to reduce the probability of underestimating the variance of individual sites and of overestimating the correlation between internal and external variables.

Call 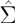 the sample estimator for Cov(*Z*). Then the 1 + λ diagonally inflated covariance is 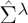 is defined as follows:

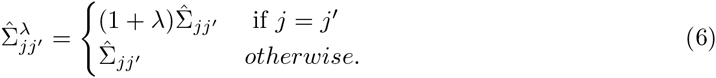

## 5 Controlling false coverage on the set of intervals

When the threshold is selected liberally, the result of running the threshold-and-merge algorithm is a large set of detected regions. If only a subset of these regions is reported, this selection could lead to low coverage properties over the selected set. In our framework, because the confidence intervals are conditioned on the initial selection, the problem does not arise (Fithian et al., 2014). However, if we use the new p-values (or intervals) to screen regions that are not separated from 0, a secondary adjustment is needed.

In that case, an iterative algorithm built described in Benjamini and Yekutieli (2005) can be applied to choose a second level inflation factor for the intervals, so that all intervals do not cover 0 and FCR is maintained. At each iteration, the set of intervals is pruned so that only intervals separated from 0 are kept in the set. Then, the BH procedure is run on the subset of p-values, selecting the q-value threshold that controls the false discovery rate at *α*. Setting new confidence intervals at the 1 – *q* level controls the rate of false coverage for the family of intervals. If any of the intervals cover 0 after the inflation, the set can be further pruned, and another BH algorithm run on the smaller set. The requirements for the usual FDR assumptions to hold over a set of regions are discussed in Schwartzman et al. (2013).

Computationally, we can reuse the sample for the p-values to fit any size of confidence interval. Here is a review of the full algorithm:

1. Run a threshold-and-cluster algorithm to generate the bump candidates.
2. For each bump, test the selective hypothesis that the effect is smaller than *ℓ* (typically, *ℓ =* 0).
3. Find the p-value for 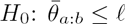.
4. Run the BH procedure on the (sorted) p-value list. Find a value q that controls the FDR at level *α.*
5. For bump candidates that pass the criterion, form marginal confidence intervals with coverage 1 − *q*. The sample from stage 2 can be reused for this purpose.

## 6 Simulation

We conducted two set of simulations to verify that the coverage properties of the confidence intervals are robust to different scenarios and to investigate power. In both simulations, data was generated from a two-group model. Detailed description of both simulations are found in Appendix C.

In the first set, we sampled repeatedly for the same region to study the coverage and power. We sampled a *D =* 5 multivariate normal vector and selected for the region *a =* 2, *b =* 4. We varied the number of samples, the covariance shapes, the true effect size, the shape of the mean vector. We used either the true covariance, or a covariance that was estimated from the data. Note that increasing the number of samples reduces the variance of the *Z* vector, as well as improves the estimation of the covariance.

In the second set, we sampled from longer (D=50) random non-stationary continuous processes, with a random non-null difference between the groups. Marginals followed either a normal or logistic-transformed normals. For each simulation, we ran the threshold-and-merge procedure and randomly selected one of the detected ROIs. We measured coverage properties for randomly selected bump locations. The variance level of the samples and threshold were selected to be similar to those in samples from DNA-methylation arrays shown in Section 7.

### Results

Representative results for both known and estimated covariances are shown in Figure 6, and the coverage rate for *C*_*cor*_ is summarized in Figure 7 (left). For the known covariance, coverage rates are approximately nominally correct under both covariance regimes, different group sizes and effect size. For the estimated covariance, coverage is less than the nominal rate, but the error decreases as the number of samples increases. Note that although for *n* = 16 with estimated Σ the two-sided coverage is almost correct, for *θ* = 0 the lower bound is too liberal. Figure 7 (right) plots the power of the selective intervals, as the probability of not covering 0 with increasing true effects *θ*. We use only the known Σ which gives accurate coverage statements.

**Figure 6:**
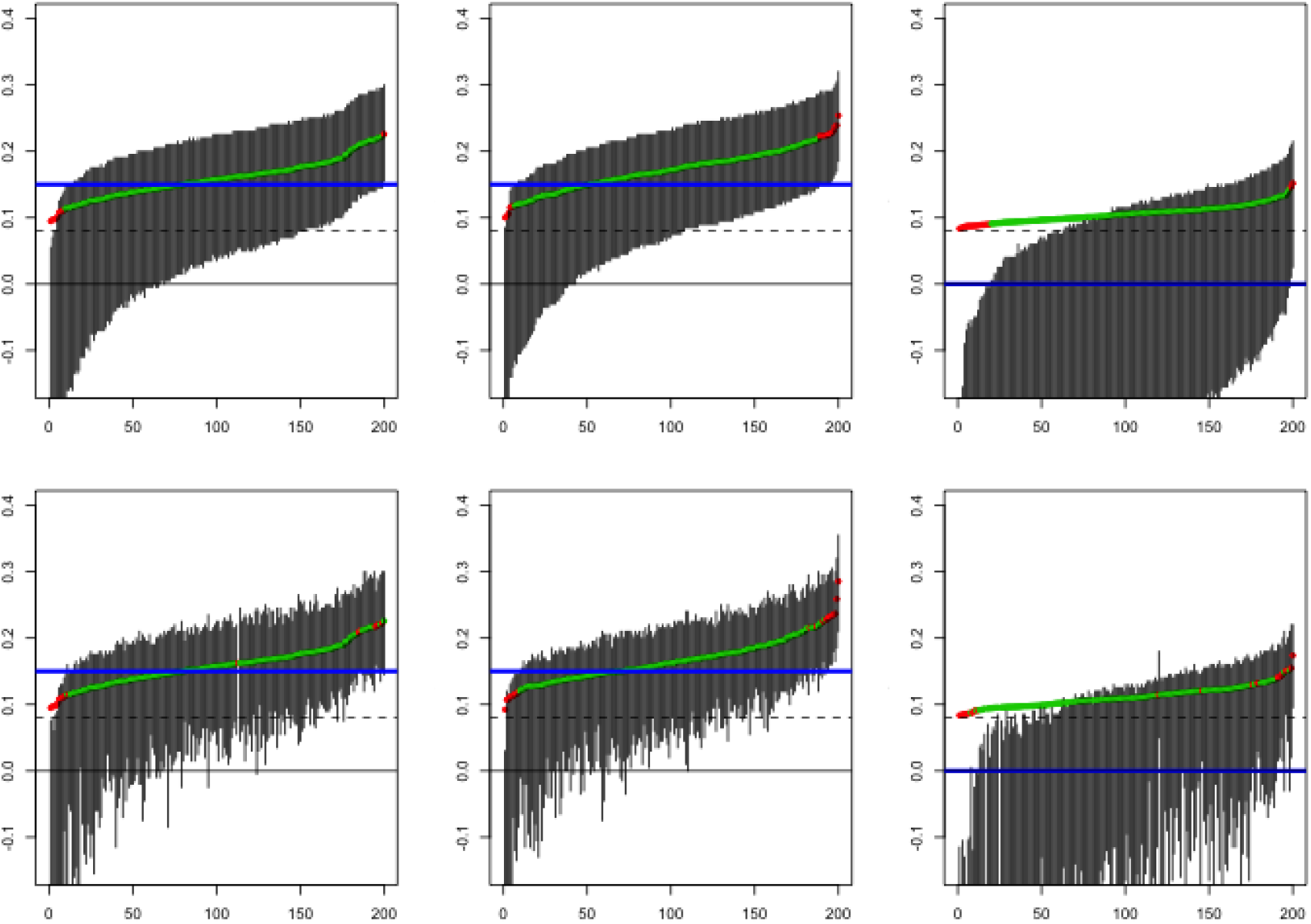
Simulations. Two sided 90% confidence intervals for repeated samples of 3-measurement bumps. Columns differ by (*θ*, Σ), from left to right (0.15, Σ_*iid*_), (0.15, *Σ*_*cor*_), (0,*Σ*_*cor*_). On the top row Σ is known, and the bottom Σ is estimated. For each regime, the runs are sorted by the observed statistic. The blue horizontal line reflects the true mean; the dot represents observed mean *t*_*obs*_. Red dots reflect instances were the true parameter was not covered.

**Figure 7:**
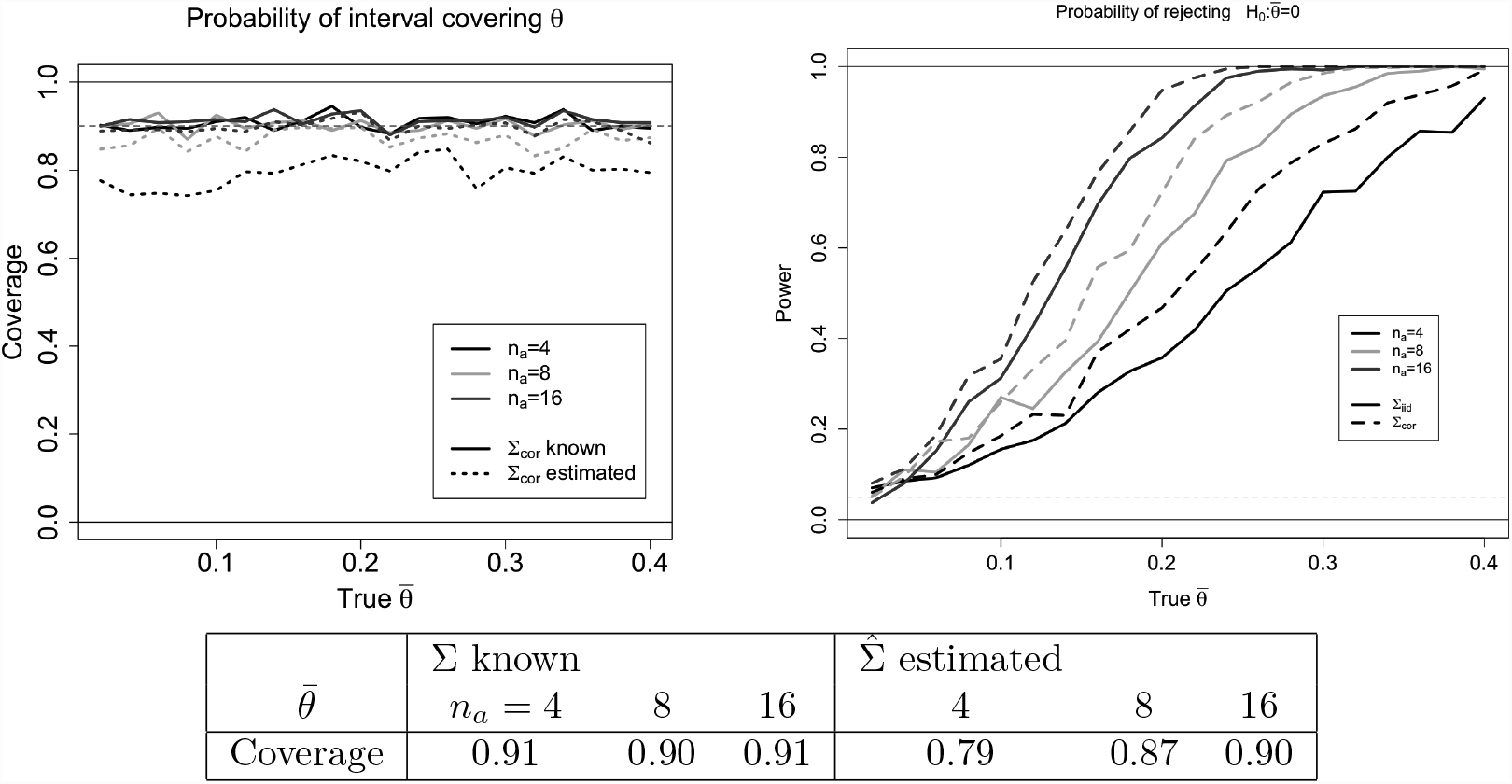
Coverage and Power: On left, coverage probability of nominal *α* = 0.9 confidence intervals for different true effect size 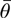 (x-axis), group size (color), and estimation of covariance (line-type). Group size affects the variance of *Z* and, if Σ is estimated, the samples available for this estimation. We see that coverage is approximately correct for the known covariance and for estimated covariance with *n*_*a*_ = 16. Note that the profile used (s_∑_) is not proportional to the true mean. On right, power is plotted for different true effect size (x-axis), group size (color), and known covariance type (*Σ*_*iid*_ or *Σ*_*cor*_). Power is computed as the proportion of intervals not covering the null for non-zero true effects. Results for estimated Σ not shown, because coverage in-exact for small *n*’s. Each value is based on 250 repeats See simulation details in Section 6. Overall average based on 5000 simulations (250 repeats × 20 values of *θ*).

The results from the continuous simulation are displayed in Table 6. For Normal data with the true covariance, we get the expected coverage of 0.9 even though the true mean vector was misspecified.

For the logistic-transformed normal data, we get conservative intervals, perhaps due to the short tails. For the estimated Σ, accuracy of coverage depends on the number of samples; increased sample size allows better estimates of Σ estimate. Still, for 10 + 10 samples the algorithm seems to give approximately correct coverage.

**Table 1:**
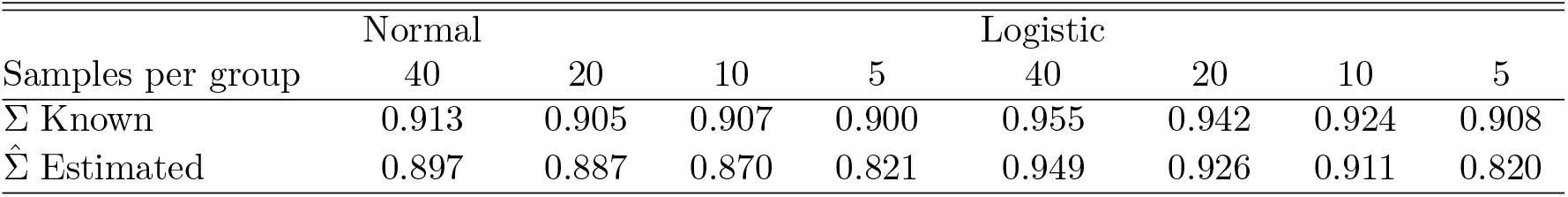
Coverage of *α* = 0.9 confidence interval for continuous process data. Values based on 1000 repeats. Variance in estimators decreases linearly with number of samples.

## 7 DNA methylation data

The inference was run on detected regions analysing healthy human samples across two different tissues. Our dataset consists of methylation measurements for 36 tissue samples using the Illumina 450K array. Data was acquired from The Cancer Genome Atlas (TCGA).

We run two analyses of the data:

- The *two-tissue* analysis compares 19 lung samples with 17 colon samples. We expect many true differences between the two groups.
- For the *one-tissue* data, we randomly partitioned the 19 lung samples into two groups of 9 and 10 samples. Group-difference found on the one-tissue data are considered false-positive.

On each dataset, we detected ROIs using a fixed threshold (*c* = 0.1) to produce a list of candidate regions. For detection, we used the *bumphunter* (*v1.10.0*) package (Irizarry et al.). For each detected ROI, we produced a selective p-value and formed a 90% selective confidence interval for mean between-group difference. The sample estimator of Σ was used for inference. Regions whose intervals overlapped 0 were pruned and intervals readjusted using (a) BH procedure to control FCR as discussed in Section 5 or (b) Bonferroni procedure to control family-wise probability of non-coverage. The samples were not smoothed or preprocessed. We did not allow regions to include sites separated by more than 5000 bps. The analysis was implemented in R. The TMN distribution was sampled using a C-compiled version of the *selective-inference* sampler, accelerated by tilting. For comparison, we produced p-values for the same set of detected ROIs using the family-wise error correction using permutations (Jaffe et al., 2012b) as implemented in *bumphunter*. All candidate regions were ranked by area, and compared to the strongest regions found in a null distribution that assumes random assignment to groups. The FWE corrected p-value was set to the proportion of permuted datasets in which a superior region was found.

### Results

The summary results for the two tissue and the one tissue data are summarized in Table 2. The selective inference method displays a much greater sensitivity than the permutation level, while giving little in terms of specificity. For the two tissue design, selective inference calls 89% of the found regions to be significantly different from 0 at the 0.05 BH adjusted level, and 63% for an FWE of 0.05. This compares to 0.07% for the permutation based FWE at 0.05. Examples for regions detected by selective inference and not by permutation FWE are shown in Figure 8.

**Table 2:**
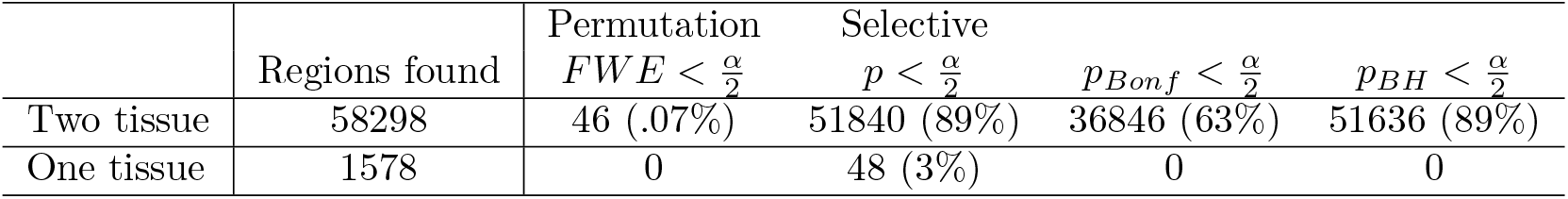
Number of regions detected on the two-tissue and one-tissue designs, for one-sided *α*/2 = 0.05 tests. Data in the one-tissue were split randomly into two groups, so we consider all detections to be false positives. Estimated covariance 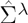 were used with *λ* = 0.15.

| | Regions found | Permutation<br>$FWE < \frac{\alpha}{2}$ | Selective<br>$p < \frac{\alpha}{2}$ | $p_{Bonf} < \frac{\alpha}{2}$ | $p_{BH} < \frac{\alpha}{2}$ |
| --- | --- | --- | --- | --- | --- |
| Two tissue | 58298 | 46 (.07%) | 51840 (89%) | 36846 (63%) | 51636 (89%) |
| One tissue | 1578 | 0 | 48 (3%) | 0 | 0 |

**Figure 8:**
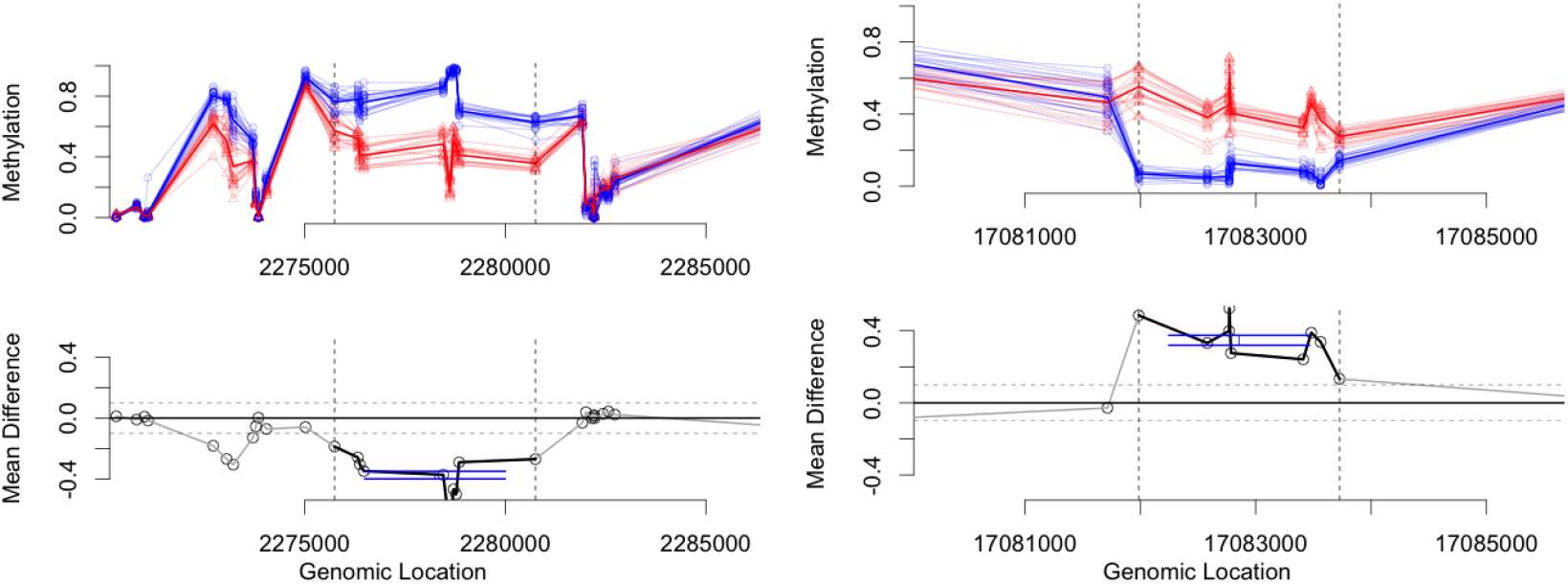
Examples undetected by non-parametrics. Two example regions that are detected using our method, but not detected using the non-parametric FWE approach at *α* = 0.05 level. On left is a 10-site region from chromosome 19: we estimate 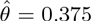 with interval 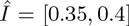 and 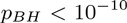; non-parametric FWE was 0.06. On right is a 9-site region from chromosome 22: we estimate 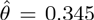 with interval 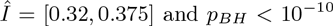; non-parametric FWE was 0.1. Data is from TCGA; see details in text.

For the one tissue design, the nominal coverage of the intervals is conservative (3% of regions are rejected at the 0.05 level). No region is significant after multiplicity corrections with either method. If no covariance inflation is used (*λ* = 0), 5.5% of the regions are rejected at the 0.05 level, and 11 regions pass the *BH* procedure.

## 8 Discussion

In this paper we present a method of generating selection corrected p-values, estimates and confidence intervals for the effect size of individual regions detected from the same data. The method allows for non-stationary individual processes, as each region is evaluated according to its own co-variance. For a two group design under non-negative correlation, the nominal coverage of the tests and lower-bounds of the intervals are shown to hold when the covariance is known or the number of samples per group is moderate (group size ≥ 16). For genomic data sets, we show that the method has considerably better power than non-parametric alternatives, and the resulting intervals are often short enough to aid decision making. In the following we discuss further

### Setting the threshold

The threshold *c* has considerable sway over the size and the number of regions detected. For the method to be effective requires a reasonable threshold: setting it too high will “condition away” all the information at the selection stage, and very little information will be used for the inference. Setting the threshold too low will allow many regions to pass, requiring stronger multiplicity corrections to control the family-wise error rates. Exploring this tradeoff via a higher-criticism (Donoho and Jin, 2008) type approach may be an interesting extension.

More generally, the assumption that *c* is determined before the analysis can probably be relaxed, as the threshold is only very weakly dependent on any individual region. In particular, data-dependent thresholds computed far from the selected region (e.g. on different chromosomes) would have similar properties; a possible algorithm is then to compute a threshold for each chromosome based on data from all other chromosomes. Robust function of the data such as the median or non-extremal quantile based thresholds should also give good results. See Weinstein et al. (2013) for an adaptive threshold in univariate selection.

### Region size information

A related question is how to integrate the information regarding the size of the region. The methods we propose do not directly use information regarding the size of the detected region, as this information is conditioned away. Instead, inference is based only on the distance between the observations and the threshold: if the observed values are sufficiently close to the threshold, the p-values will be large; if they are farther away, p-values will be small. Discarding the size of the region is perhaps counterintuitive. Hypothetically, we may detect a large enough region (with *P*(*A*_*a:b*_) small enough) to be significant regardless of the effects of selection, but still get selective p-value that are large. In practice, however, this is unlikely; both the probability of the event *A*_*a:b*_ and the selective p-value become smaller as the size of the unconditional mean vector (Θ_*a:b*_) increases. Long regions would usually be detected because the mean was larger than 0 in most of the region. This would usually also manifest in smaller p-values and less uncertainty in the confidence interval.

Reintegrating the probability of detection can increase power, but would require a different mechanism to control for selection. We may be able to recover the probability of selection from the Monte Carlo sample. It is tempting to reintegrate this information into the inference: under a strong null (Θ_*a:b*_ = 0), the likelihood of the data is the product of these two probabilities. The caveat, of course, is that the p-values associated with region size – *P*(*A*_*a:b*_) – are not corrected for selection. Hence, we are back to the problem we wanted to initially solve.

### Difference between our approach and pivot-based methods

The methods we propose are different from the exact pivot-based inference advocated by, for example,? and?. These advocate conditioning not only on the selection event, but also on the subspace orthogonal to the statistic of interest. Essentially, all but the one-dimension statistic *t*(*Z*) are conditioned away, leaving a well-specified single-parameter conditional distribution to evaluate. The model is elegant, in that it produces an exact p-value and intervals, without requiring sampling or plugging in nuisance parameters. However, we found that the fully-conditional approach had very little power separate true effects from nulls, and resulted in very large intervals. Specifically, after the conditioning, inference is conducted within a single segment {*Z*_*a:b*_ *+ α*(1,…, 1)}_*α*_; if the estimate of any of the points in the region is very close to the threshold, the will be no separation and the p-value obtained would be high. In contrast, our method is not sensitive to having individual points which are close to the threshold, because it aggregates outcomes over the set 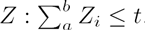. We pay a price in having an inexact method that leans on sampling and a misspecified choice of mean vector.

### Additional applications and future work

The importance of accurate regional inference inference extends from only genomics. Indeed, the threshold-and-merge region detection algorithm is extremely common in neuroscience for the analysis of fMRI data, where it is known as cluster inference. The standard parametric methods used for cluster inference rely on approximations for extreme sets in stationary gaussian processes (Friston et al., 1994). Recently, the high-profile study of (Eklund et al., 2016) showed these methods to be too liberal by testing them on manufactured null; also, the distribution of detections was not uniform along the brain suggesting the process was non-stationary. The alternative offered were non-parametric permutations of subject assignment (Hayasaka et al., 2004), similar to those used in Section 7. As we observe, these non-parametric methods can be grossly over-conservative in their model, in particular when multiple regions are detected. Adapting our method for functional data may allow a powerful parametric model that relaxes the stationarity and strong thresholding requirements, without sacrificing power.

More work is required to expand the scope of the algorithm for these additional applications. In particular, the method needs to work well for larger regions and smaller samples. Currently, for larger regions, sampling becomes harder due to increased sensitivity of the initial parameter and to the mixing time of the Gibbs sampler. Solutions for these problems include more robust sampling algorithms and convergence decisions. Perhaps, for larger regions, approximations of *t*(*Z*)|*A* can replace Monte Carlo methods.

Furthermore, local covariance estimates might require too many samples to stabilize, and rigorous methods should be employed to deal with the unknown covariances. Using the truncated multivariate t instead of multivariate normal would account for the uncertainty in estimating the variances; however, the correlation structure also has uncertainty which we currently do not take into account. We suggest in (6) an inflation parameter to give a conservative estimate of the correlation, leaving to the user the choice of *λ*. It is more likely that for each application, specific models for covariance estimation can be developed. In genomics, external annotation including probe-distance and sequence composition can give a prior model for shrinkage.

## 9 Acknowledgements

The authors would like to acknowledge Will Fithian, Yossi Rinott and Yoav Benjamini for their comments that contributed to this manuscript. Y.B. would also want to thank Giovanni Parmigiani and the Stein Fellowship at Stanford for sponsoring a summer stay at the Biostatistic Department at Dana Farber that initiated this research. R.A.I is supported by NIH grant R01GM083084.

## A Appendix: Accelerated sampling

Recall that we parameterize the truncated multivariate normal family with a single mean parameter *θ* that linearly determines the mean vector. The distributions corresponding to different values of *θ* form a single parameter exponential family; this implies that importance weighting can efficiently convert a sample for 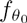 into a sample for 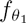. We detail here the algorithm. This is an expansion of the ideas described in the appendix of (Fithian et al., 2014).

Suppose we have a Monte Carlo sample 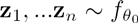 and we would like to estimate of **E**[*g*(*Z*)] for 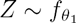. Then the importance sampling estimate of 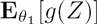 is

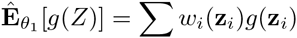

where

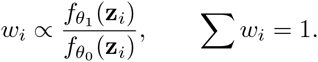

The importance estimator is unbiased. With a careful choice of *θ*_0_ it may enjoy lower variance per sample-size compared to Monte Carlo estimates from 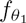 (Owen, 2013). Nevertheless, for our application the primary gain is the ability to invert tests for numerous values of *θ* using a single sample.

The exponential tilting principle (Siegmund, 1976) recognizes that for single-parameter exponential families, the importance weights *w*_1_,…,*w*_*n*_ can be calculated without explicitly calculating the normalizing constant for the destination density 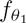. We develop here the explicit form for the TMN densities parameterized by a linear mean shift.

The TMN density (5) is written in full as:

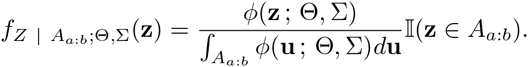

with the mean vector parameterized linearly in *θ*

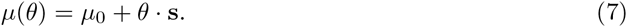

As described above, we take

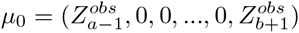

and a profile 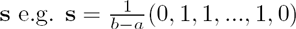.

We can expand this density to exponential family form:

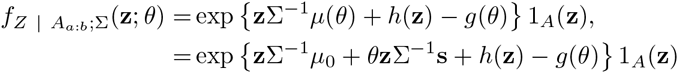

where 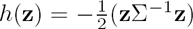 does not depend on *θ,* and 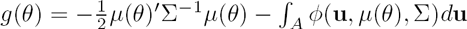 is the normalizing constant. Therefore, the likelihood ratio for an example z ∈ *A* depends on the covariance-corrected shape

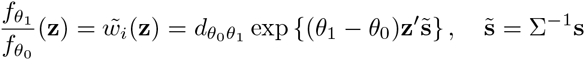

where the factor 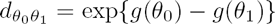 does not depend on **z**. Instead of computing 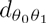 the weights for a given Monte Carlo sample 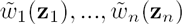 can be normalized

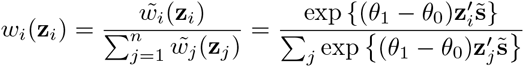

to meet both conditions of empirical importance sampling weights.

**Figure 9:**
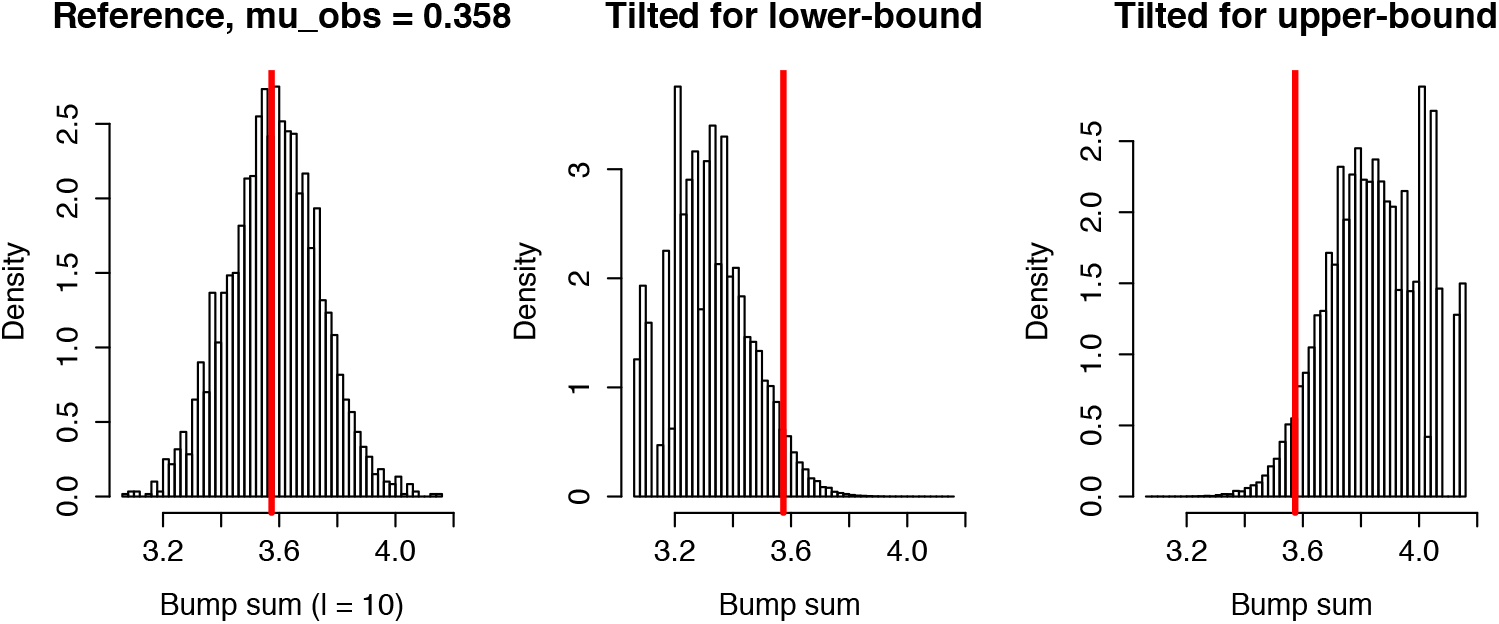
Example of how to extract confidence intervals from a reference density. We generate a sample from a reference density (and mean parameter *θ*) for which the observed statistic *t*_*obs*_ = 0.358 (red vertical line) is in the bulk of the distribution. The sample can then be tilted - reweighed - to generate other distributions within the same parametric family. The tilted distribution giving the lower bound of the confidence interval is plotted in the center. This is the most left-tilted distribution for which *P*(*T* > *tobs*) ≥ *α*/2. The distribution for the upper-bound is plotted on the right.

### A.1 Tilting for tests and intervals

The main speed-up in tilting arrises from the ability to use a single sample from 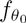 to identify the acceptance region for any *θ*. This is particularly useful because our method for estimating confidence intervals requires building many individual tests for a dense grid of *θ* values. The main computational load comes from sampling the TMN under the constraint; tilting allows us to perform this task only once, for a convenient value of *θ,* and extract all tests from this sample.

Computing the p-value for the null hypothesis *H*_0_ : *θ* ≤ 0 against a one-sided alternative *H*_1_ : *θ* > 0. To estimate the probability under the null of getting a value greater than *t*_*obs*_, or

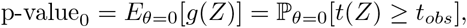

we use the quantile of the ‘tilted’ empirical distribution

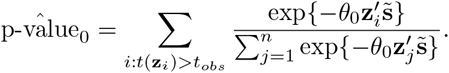

For a two sided test of *H*_0_ : *θ* = *θ*_1_, as required for two-sided intervals, we use 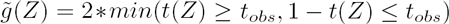. The test rejects 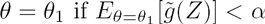.

Note that it is not necessary that the statistic *t*(*z*) coincide with the sufficient statistic 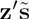 of the exponential family. Although the sufficient statistic introduces the most powerful tests, other linear statistics may be more robust to deviations from the mean model. In the case were a different statistic is preferred, the reweighing of the Monte Carlo samples might not be completely correlated to the statistic.

### A.2 Choosing *θ*_0_

Heuristically, we prefer a *θ*_0_ for which *t*_*obs*_ is a likely outcome, with enough samples on both sides of *t*_*obs*_. In the extreme case, to identify any p-value in (0,1) via tilting requires that 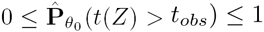, meaning we have at least one sample at each side of *t*_*obs*_. The variance of an importance sample is 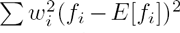 (Owen, 2013, ch 9, pg 9), so a more equal distribution of weights would lead to better variances., a likely choice for *θ*_0_ would often mean the selection criterion is not very strong, so that the samplers for *θ*_0_ would be relatively efficient.

For choosing *θ*_0_ under a truncation at *c*, one candidate choice is to use the unbiased estimator if there were no selection *θ*_0_ = *t*_*obs*_. We require the empirical 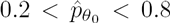. If this value fails we run a linear search for *θ*_0_ values between *t*_*obs*_ and 0 until we find a successful value. Due to selection against small values of Z, *t*_*obs*_ is an upward biased estimator of *θ*; when variances are small, the sampler might need a better starting point for *θ*_0_, so multiple starting points can be explored.

## B Appendix: Proofs and Examples

**Lemma 1**

Let 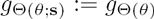 denote the family densities for *t*(*Z*) with a scale single parameter *θ*. Then

1. 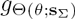 is a monotone likelihood ratio family
2. *E*[*t*(*Z*)] is an increasing function of *θ*
3. The confidence set for *θ* obtained by inverting two sided tests is an interval.

**Proof**

We recall the exponential family form of 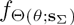:

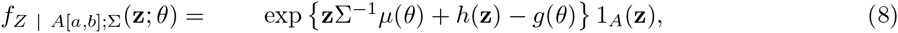

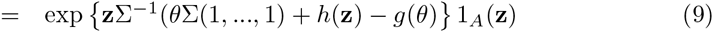

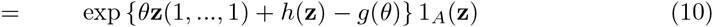

where 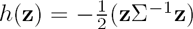 does not depend on *θ*, and 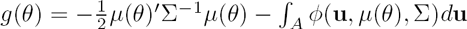 is the normalizing constant.

Next note that the density in 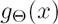 for a particular value corresponds to integrating over the intersection of the hyper-plane ∑ *Z* = *z* with the conditional set *A*. Within this hyperplane, the value of the statistic is fixed zʹ(1,…, 1) *= x* so

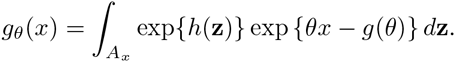

Because the expression exp{*θx* – *g*(*θ*)} is fixed within *A*_*x*_, it can be moved outside the integral. Defining

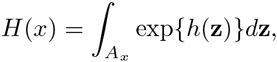

we get the exponential family structure in

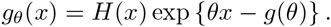

Therefore, the likelihood ratio for a value *t*(*z*) *= x* can be written as

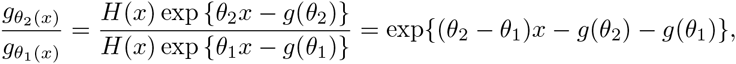

a strictly increasing function of *x.*

**Lemma 2**

We use results from Rinott and Scarsini (2006) for multivariate normals to identify the following conditions on Σ and s:

1. Σ^−1^ is an M-matrix, meaning all off-diagonal elements are non-positive. (In particular, Σ must be non-negative).
2. s lies in the cone *C*_Σ_ of non-negative linear combinations of columns of Σ

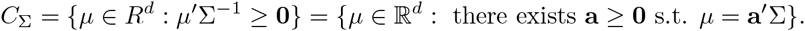

Note that for Σ = *I*, any non-negative profile would be in *C*_Σ_ (indeed, we see that all profiles produce a monotone curve for an iid covariance, see Figure 5). The condition is sufficient, but not necessary; even if Σ is not an M-matrix, there might still be specific (profile, statistic) pairs for which monotonicity will hold, as discussed in Lemma 1. However, there will be no profile for which any statistic will be monotone.

Let 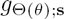 be the family of densities of *t*(*Z*)|*A* parametrized by *θ,* where *Z* ~ *N*(*θ ·* s, Σ). If Σ^−1^ is an M-matrix, and the profile s can be written as a non-negative sum of columns of Σ, then 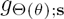 is a monotone likelihood ratio family.

**Proof**

Consider random vectors *Z* and *Z′* where *Z* ~ *N*(*θ ·* s, Σ) and *Z′* ~ *N*(*θ′* · s, Σ) for some *θ′* > *θ*. The lemma identifies a sufficient condition for *Z′|A* being stochastically larger (≥_*st*_) than *Z*|*A* Stochastic ordering implies that for any positive functional 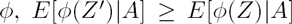, and in particular for a positive statistic 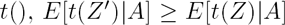 and the quantile functions of *t*(*Z*)|*A* are similarly ordered.

The key to the proof is moving from an ordering of *Z* and *Z′* into an ordering of the conditional vectors [*Z*|*A*] and [*Z*′|*A*], (interpreted as [*Z*|*Z* ∈ *A*] and 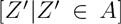). For rectangular sets *A,* the stronger notion of ordering *total positivity* is maintained through the conditioning. We review here the main results of Rinott and Scarsini (2006); proofs and an extended discussion of the cited properties are found there.

- For multivariate densities *f*, *g,* the relation 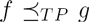 (total positivity) implies that 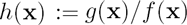 is increasing in any coordinate-wise increase in x. [Lemma 2.2, a]
- If 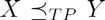, then *X* is also stochastically greater than *Y*, implying 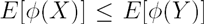 for any nondecreasing function *ϕ* [Proposition 2.4]. If 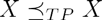 then *X* is said to be *multivariate total positive of order 2* or *MTP2*.
- For a rectangular set 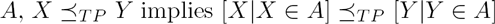 [Theorem 2.5, a special condition of Remark 2.6 (i)] Note that conditioning by thresholding individual coordinates would always result in a rectangular set *A.*
- A multivariate normal *Z* with an invertible covariance matrix Σ is MTP2 if and only if Σ^−1^ is an M-matrix. [2.15]
- If for some 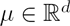 we have 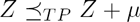, then *Z* is *MTP*_2_. [Thm 3.2]
- For *Z* that is *MTP*_2_, we have 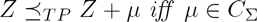 [Thm 3.2]

Therefore, under the assumptions regarding Σ, *Z* is *MTP –* 2. Call 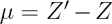, then *µ* = (*θ′ – θ*) · *s* so 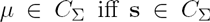. Therefore, the conditions suffice for 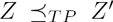. This further implies 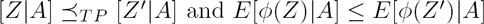.

## C Appendix: Simulation Details

Details of the two simulation studies are described below.

**Experiment 1**

In the first set of experiments, we tested coverage probability of 1 – 2*α* = 0.9 confidence intervals by repeatedly sampling the same set of variables, keeping only those data-vectors for which all locations passed the selection threshold. We sample data vectors of length *D =* 5, selecting for the *positive* region *r* = (2, 4, +) spanning the vector. Data was generated from the multivariate normal with mean bump-heights of 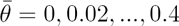. For the true mean vector we used 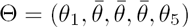 with 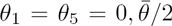. Sample size was *n*_1_ = *n*_2_ = 4, 8,16; increased sample size reduces the variance of *Z* and, for unknown covariance, increases the accuracy of 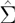. The data was generated using two covariance matrices for the samples: a correlated *C*_*cor*_ and an uncorrelated *C*_*iid*_,

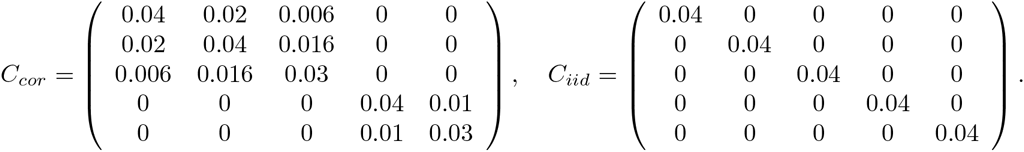

In both matrices, *σ*^2^ = 0.04 was chosen to reflect the average observed within-group variance in the DNA-methylation data.

The following results are based on a tilting algorithm using *n =* 12000 samples from the reference distribution. The confidence interval estimation was repeated *N =* 2000 times for 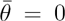 and *N =* 250 each value 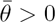. We repeated each experiment three times, using (a) the true covariance, (b) the sample covariance 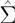, and (c) the inflated covariance 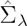 with λ = 0.15, see (6). We used the profile 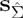 in all experiments: under the known iid covariance, this corresponds to the true shape of the mean; in other cases the profile doesn’t match the shape of the true mean.

**Experiment 2**

Data was generated in the following way:

1. The (untransformed) mean process for group A *µ*_*A*_ was generated by sampling D=50 data points from an iid *N*(0,1) process, and convolving with the 1-dimensional kernel *K*_*µ*_ *=* (0.1, 0.2,0.4, 0.2,0.1). The mean difference process *µ*_*δ*_ was generated by sampling iid 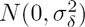 and convolving with *K*_1_. The mean for group 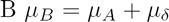.
2. For each sample we added noise 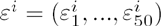 by concatenating noise from two correlation regimes:
  - For 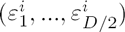, iid *N*(0,1) samples were smoothed by convolving each vector with *K*_*ε*_ = (0.05,0.1,0.15,0.4,0.15,0.1,0.05). This resulted in correlated noise with 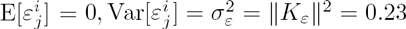.
  - For 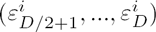, iid noise was sampled from 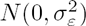 with no smoothing.
3. Noise was added to each sample so that 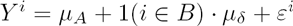.
4. For the transformed data, each sample *Y*^*i*^ was transformed coordinate-wise with the logistic function 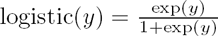. For the transformed data, a population of *N* = 10000 samples was generated, and the mean vector and covariance matrix for each group were estimated empirically from the samples.
5. Multiple subsamples of *n =* 40, 20,10, 5 were taken from each group.

## Footnotes

1 Recall that *p* is the number of sample covariates – *length*{(*X*_*i*_, *W*_*i*_)} – which is typically small, not to be confused with the size of the measurement vector *D* = *length*{*Y*_*i*_}.

2 Admittedly, we reverse here the usual flow of statistical modeling, by first choosing the statistic and only later the model.

## References

MJ Aryee, AE Jaffe, H Corrada-Bravo, C Ladd-Acosta, AP Feinberg, KD Hansen?, and RA Irizarry. Minfi: A flexible and comprehensive bioconductor package for the analysis of infinium dna methylation microarrays. Bioinformatics, 30(10):1363–1369, 2014.

Claude Becker, Jörg Hagmann, Jonas Müller, Daniel Koenig, Oliver Stegle, Karsten Borgwardt, and Detlef Weigel. Spontaneous epigenetic variation in the arabidopsis thaliana methylome. Nature, 480(7376):245–249, 2011.

Yoav Benjamini and Daniel Yekutieli. False discovery rate–adjusted multiple confidence intervals for selected parameters. Journal of the American Statistical Association, 100(469):71–81, 2005.

Yuval Benjamini and Terence P. Speed. Summarizing and correcting the gc content bias in high-throughput sequencing. Nucleic Acids Research, 40(10):e72, 2012.

Richard Berk, Lawrence Brown, Andreas Buja, Kai Zhang, Linda Zhao, et al. Valid post-selection inference. The Annals of Statistics, 41(2):802–837, 2013.

Marina Bibikova, Bret Barnes, Chan Tsan, Vincent Ho, Brandy Klotzle, Jennie M Le, David Delano, Lu Zhang, Gary P Schroth, Kevin L Gunderson, et al. High density dna methylation array with single cpg site resolution. Genomics, 98(4):288–295, 2011.

Christoph Bock, Jörn Walter, Martina Paulsen, and Thomas Lengauer. Inter-individual variation of dna methylation and its implications for large-scale epigenome mapping. Nucleic Acids Research, 36(10):e55, 2008. doi:10.1093/nar/gkn122. URL http://nar.oxfordjournals.org/content/36/10/e55.abstract.

T Tony Cai and Ming Yuan. Rate-optimal detection of very short signal segments. arXiv preprint arXiv:1407.2812, 2014.

Thomas J DiCiccio and Joseph P Romano. On parametric bootstrap procedures for second-order accurate confidence limits. Technical report, Technical report, 1988.

David Donoho and Jiashun Jin. Higher criticism thresholding: Optimal feature selection when useful features are rare and weak. Proceedings of the National Academy of Sciences, 105(39): 14790–14795, 2008.

Bradley Efron. Bootstrap confidence intervals for a class of parametric problems. Biometrika, 72 (1):45–58, 1985.

Anders Eklund, Thomas E Nichols, and Hans Knutsson. Cluster failure: Why fmri inferences for spatial extent have inflated false-positive rates. Proceedings of the National Academy of Sciences, page 201602413, 2016.

Andrew P Feinberg and Benjamin Tycko. The history of cancer epigenetics. Nature Reviews Cancer, 4(2):143–53, Feb 2004. doi:10.1038/nrc1279.

William Fithian, Dennis Sun, and Jonathan Taylor. Optimal Inference After Model Selection. ArXiv e-prints, -, October 2014.

Karl J Friston, Keith J Worsley, RSJ Frackowiak, John C Mazziotta, and Alan C Evans. Assessing the significance of focal activations using their spatial extent. Human brain mapping, 1(3):210–220, 1994.

John Geweke. Efficient simulation from the multivariate normal and student-t distributions subject to linear constraints and the evaluation of constraint probabilities. Citeseer, 1991.

Donald J Hagler, Ayse Pinar Saygin, and Martin I Sereno. Smoothing and cluster thresholding for cortical surface-based group analysis of fmri data. Neuroimage, 33(4):1093–1103, 2006.

KD Hansen, B Langmead, and RA Irizarry. Bsmooth: from whole genome bisulfite sequencing reads to differentially methylated regions. Genome Biology, 13(10):R83, 2012.

Satoru Hayasaka and Thomas E. Nichols. Validating cluster size inference: random field and permutation methods. NeuroImage, 20(4):2343–2356, 2003. ISSN 1053-8119. doi:http://dx.doi.org/10.1016/j.neuroimage.2003.08.003. URL http://www.sciencedirect.com/science/article/pii/S1053811903005020.

Satoru Hayasaka, K. Luan Phan, Israel Liberzon, Keith J. Worsley, and Thomas E. Nichols. Non-stationary cluster-size inference with random field and permutation methods. NeuroImage, 22 (2):676–687, 2004. ISSN 1053-8119. doi:http://dx.doi.org/10.1016/j.neuroimage.2004.01.041. URL http://www.sciencedirect.com/science/article/pii/S1053811904000862.

William C Horrace. Some results on the multivariate truncated normal distribution. Journal of Multivariate Analysis, 94(1):209–221, 2005.

Rafael A Irizarry, Martin Ayree, Kasper D Hansen, and Hector C Hansen. bumphunter: Bump hunter. URL https://github.com/ririzarr/bumphunter.

Rudolf Jaenisch and Adrian Bird. Epigenetic regulation of gene expression: how the genome integrates intrinsic and environmental signals. Nature genetics, 33:245–254, 2003.

Andrew E Jaffe, Andrew P Feinberg, Rafael A Irizarry, and Jeffrey T Leek. Significance analysis and statistical dissection of variably methylated regions. Biostatistics, 13(1):166–178, 2012a.

Andrew E Jaffe, Peter Murakami, Hwajin Lee, Jeffrey T Leek, M Daniele Fallin, Andrew P Feinberg, and Rafael A Irizarry. Bump hunting to identify differentially methylated regions in epigenetic epidemiology studies. International Journal of Epidemiology, 41(1):200–209, 2012b. doi:10.1093/ije/dyr238. URL http://ije.oxfordjournals.org/content/41/1/200.abstract.

Andrew E Jaffe, Peter Murakami, Hwajin Lee, Jeffrey T Leek, M Daniele Fallin, Andrew P Feinberg, and Rafael A Irizarry. Bump hunting to identify differentially methylated regions in epigenetic epidemiology studies. International journal of epidemiology, 41(1):200–209, 2012c.

Nikolaus Kriegeskorte, W Kyle Simmons, Patrick SF Bellgowan, and Chris I Baker. Circular analysis in systems neuroscience: the dangers of double dipping. Nature neuroscience, 12(5): 535–540, 2009.

Pei Fen Kuan and Derek Y. Chiang. Integrating prior knowledge in multiple testing under dependence with applications to detecting differential dna methylation. Biometrics, 68(3):774–783, 2012. ISSN 1541-0420. doi:10.1111/j.1541-0420.2011.01730.x. URL http://dx.doi.org/10.1111/j.1541-0420.2011.01730.x.

Pei Fen Kuan, Sijian Wang, Xin Zhou, and Haitao Chu. A statistical framework for illumina dna methylation arrays. Bioinformatics, 26(22):2849–2855, 2010. doi:10.1093/bioinformatics/btq553. URL http://bioinformatics.oxfordjournals.org/content/26/22/2849.abstract.

Anshul Kundaje, Wouter Meuleman, Jason Ernst, Misha Bilenky, Angela Yen, Alireza Heravi-Moussavi, Pouya Kheradpour, Zhizhuo Zhang, Jianrong Wang, Michael J Ziller, et al. Integrative analysis of 111 reference human epigenomes. Nature, 518(7539):317–330, 2015.

Jason D. Lee, Dennis L. Sun, Yuekai Sun, and Jonathan Taylor. Exact post-selection inference with the lasso. ArXiv e-prints, November 2013.

Lung-Fei Lee. Consistent estimation of a multivariate doubly truncated or censored tobit model. Discussion Paper No. 153, 1981.

Jeffrey T Leek, Robert B Scharpf, H´ector Corrada Bravo, David Simcha, Benjamin Langmead, W Evan Johnson, Donald Geman, Keith Baggerly, and Rafael A Irizarry. Tackling the widespread and critical impact of batch effects in high-throughput data. Nature Reviews Genetics, 11(10): 733–739, 2010.

E. L. Lehmann and J. P. Romano. Testing Statistical Hypotheses (Third Edition). Springer, 2005.

Ryan Lister, Mattia Pelizzola, Robert H Dowen, R David Hawkins, Gary Hon, Julian Tonti-Filippini, Joseph R Nery, Leonard Lee, Zhen Ye, Que-Minh Ngo, et al. Human dna methylomes at base resolution show widespread epigenomic differences. nature, 462(7271):315–322, 2009.

Ryan Lister, Eran A Mukamel, Joseph R Nery, Mark Urich, Clare A Puddifoot, Nicholas D Johnson, Jacinta Lucero, Yun Huang, Andrew J Dwork, Matthew D Schultz, et al. Global epigenomic reconfiguration during mammalian brain development. Science, 341(6146):1237905, 2013.

Richard Lockhart, Jonathan Taylor, Ryan J. Tibshirani, and Robert Tibshirani. A significance test for the lasso. The Annals of Statistics, 42(2):413–468, 04 2014. doi:10.1214/13-AOS1175. URL http://dx.doi.org/10.1214/13-AOS1175.

Art B. Owen. Monte Carlo theory, methods and examples. 2013.

Alain Pacis, Ludovic Tailleux, Alexander M Morin, John Lambourne, Julia L MacIsaac, Vania Yotova, Anne Dumaine, Anne Danckaert, Francesca Luca, Jean-Christophe Grenier, et al. Bacterial infection remodels the dna methylation landscape of human dendritic cells. Genome research, 2015.

Ari Pakman and Liam Paninski. Exact hamiltonian monte carlo for truncated multivariate gaussians. Journal of Computational and Graphical Statistics, 23(2):518–542, 2014.

Brent S. Pedersen, David A. Schwartz, Ivana V. Yang, and Katerina J. Kechris. Comb-p: software for combining, analyzing, grouping and correcting spatially correlated p-values. Bioinformatics, 28(22):2986–2988, 2012. doi:10.1093/bioinformatics/bts545. URL http://bioinformatics.oxfordjournals.org/content/28/22/2986.abstract.

Aharon Razin and Arthur D Riggs. Dna methylation and gene function. Science, 210(4470): 604–610, 1980.

Yosef Rinott and Marco Scarsini. Total positivity order and the normal distribution. Journal of Multivariate Analysis, 97(5):1251–1261, 2006.

Keith D Robertson. Dna methylation and human disease. Nature Reviews Genetics, 6(8):597–610, 2005.

Armin Schwartzman, Yulia Gavrilov, and Robert J. Adler. Multiple testing of local maxima for detection of peaks in 1d. The Annals of Statistics, 39(6):3290–3319, 12 2011. doi:10.1214/11-AOS943. URL http://dx.doi.org/10.1214/11-AOS943.

Armin Schwartzman, Andrew Jaffe, Yulia Gavrilov, and Clifford A. Meyer. Multiple testing of local maxima for detection of peaks in chip-seq data. The Annals of Applied Statistics, 7(1):471–494, 03 2013. doi:10.1214/12-AOAS594. URL http://dx.doi.org/10.1214/12-AOAS594.

Jonathan Sebat, B Lakshmi, Jennifer Troge, Joan Alexander, Janet Young, Pär Lundin, Susanne Månér, Hillary Massa, Megan Walker, Maoyen Chi, et al. Large-scale copy number polymorphism in the human genome. Science, 305(5683):525–528, 2004.

David Siegmund. Importance sampling in the monte carlo study of sequential tests. The Annals of Statistics, pages 673–684, 1976.

David Siegmund, Benjamin Yakir, and Nancy Zhang. The false discovery rate for scan statistics. Biometrika, 98:979–985, 2011.

Max Sommerfeld, Stephen Sain, and Armin Schwartzman. Confidence regions for excursion sets in asymptotically gaussian random fields, with an application to climate. arXiv preprint arXiv:1501.07000, 2015.

Lingyun Song and Gregory E Crawford. Dnase-seq: a high-resolution technique for mapping active gene regulatory elements across the genome from mammalian cells. Cold Spring Harbor Protocols, 2010(2):pdb–prot5384, 2010.

Wenguang Sun, Brian J Reich, T Tony Cai, Michele Guindani, and Armin Schwartzman. False discovery control in large-scale spatial multiple testing. Journal of the Royal Statistical Society: Series B (Statistical Methodology), 2014.

Daiya Takai and Peter A Jones. Comprehensive analysis of cpg islands in human chromosomes 21 and 22. Proceedings of the national academy of sciences, 99(6):3740–3745, 2002.

Asaf Weinstein, William Fithian, and Yoav Benjamini. Selection adjusted confidence intervals with more power to determine the sign. Journal of the American Statistical Association, 108(501): 165–176, 2013.

Choong-Wan Woo, Anjali Krishnan, and Tor D Wager. Cluster-extent based thresholding in fmri analyses: pitfalls and recommendations. Neuroimage, 91:412–419, 2014.

Nancy R Zhang and David O Siegmund. Model selection for high-dimensional, multi-sequence change-point problems. Statistica Sinica, pages 1507–1538, 2012.

Yong Zhang, Tao Liu, Clifford A Meyer, J´erˆome Eeckhoute, David S Johnson, Bradley E Bernstein, Chad Nusbaum, Richard M Myers, Myles Brown, Wei Li, et al. Model-based analysis of chip-seq (macs). Genome biology, 9(9):1, 2008.

